# Therapeutic interleukin-2 rewires skeletal-immune circuits to reverse postmenopausal bone loss

**DOI:** 10.64898/2026.01.07.696511

**Authors:** Pengcheng Zhou, Jinhong Feng, Yifan Zhang, Yuhua Liu, Dongchao Ma, Anqi Xue, Wei Wei, Lei Sun, Di Yu, Ting Zheng

## Abstract

Postmenopausal osteoporosis results from excessive osteoclast activity, but the immune circuits that limit osteoclastogenesis in the bone marrow remain poorly understood. Here, we tested a clinically accessible immunotherapy low-dose interleukin-2 (IL-2) as a therapeutic intervention in mice with established ovariectomy-induced bone loss, integrating microcomputed tomography imaging, single-cell transcriptomics, genetic perturbation, and human monocyte-osteoclast assays. Therapeutic administration of IL-2 reversed trabecular bone loss, expanded CXCR3⁺ regulatory T cells in bone marrow, and promoted IL-10- and CGRP-producing ILC2s that were required for protection. IL-2 also acted directly on the osteoclast lineage, suppressing mouse and human osteoclastogenesis through activation of a STAT1-IRF1-dependent IFN program in RANKL-stimulated monocytes. Together, these data define an IL-2-responsive skeletal-immune network that can be therapeutically engaged to suppress osteoclastogenesis, which opens new avenues for cytokine-based immunotherapies in postmenopausal osteoporosis and related bone disorders.

## INTRODUCTION

Postmenopausal osteoporosis is one of the major global health challenges, characterized by systemic bone loss and increased susceptibility to severe fractures (*1, 2*). The significant decline in estrogen during and after menopause disrupts bone remodeling by increasing bone resorption over bone formation (*3*). Although current antiresorptive and anabolic therapies are effective, they are often limited by long-term side effects (*4*), plateauing efficacy, or rapid bone loss upon discontinuation (*5, 6*). Emerging evidence suggests that the etiology of postmenopausal bone loss is fundamentally influenced by dysregulated immune microenvironment that fuels osteoclastogenesis. Consequently, targeting the upstream immune circuits that govern bone resorption offers a promising, yet underexploited therapeutic avenue.

Interleukin-2 (IL-2) is a pleiotropic cytokine essential for immune homeostasis (*7*). Efforts to harness IL-2 for therapeutic purposes have been underway for decades, initially focused on expanding cytotoxic lymphocytes for cancer immunotherapy (*8*), which led to U.S. Food and Drug Administration (FDA) approval of clinical use of recombinant IL-2 in human in 1992. More recently, low-dose IL-2 therapy (< 100 million I.U per day) has emerged as a promising treatment for a broad range of autoimmune and inflammatory diseases in patients, including systemic lupus erythematosus (SLE) (*9–14*), vasculitis (*15*), chronic graft-versus-host (GVHD) disease (*16*), and type 1 diabetes (*17*). A central mechanism underlying these clinical benefits is the selective expansion of regulatory T cells (T_REG_), which restores immune balance and improves disease pathology (*18–21*). In the skeletal system, IL-2-deficient mice spontaneously develop osteopenia (*22*), implicating IL-2 as a critical physiological regulator of bone density. Moreover, CD25^+^ Foxp3^+^ T_REG_ cells have been shown to directly suppress osteoclast differentiation *in vitro* (*23, 24*). However, whether therapeutic IL-2 promotes a specialized T_REG_ population that are enriched in bone marrow and thereby reverse established bone loss is not known.

In addition to T_REG_ cells, group 2 innate lymphoid cells (ILC2s) are emerging as an important IL-2-responsive population *in vivo* (*25, 26*). This is particularly relevant in immunodeficiency related to T_REG_ cells or Tregopathy (*27*), wherein T_REG_ cells are inadequate to respond to IL-2 therapy and are unlikely to confer benefit. ILC2s, through the constitutive expression of the high-affinity IL-2 receptor α (*Il2ra*, CD25), may respond robustly to IL-2 in this context. Moreover, ILC2s produce canonical type 2 cytokines, including IL-4, IL-5 and IL-13, as well as IL-10 and neuropeptides, enabling them to exert regulatory effects and promote tissue repair (*28–32*). Although ILC2s are well characterized in barrier tissues, their contribution to skeletal-immune regulation remains poorly understood. Whether IL-2 treatment can potentiate ILC2s to improve osteoporotic bone loss has not been defined.

Monocytes and macrophages express IL-2 receptors (*33–35*), but how IL-2 signaling influences their differentiation toward osteoclasts is poorly understood. Engagement of the IL-2 receptor activates JAK-STAT pathways (*36*), including STAT1 in immune cells that can induce downstream transcriptional programs such as IRF1 (*37*). Notably, IRF1-dependent interferon pathways are known to respond to cytokines (*38*) and constrain osteoclastogenesis (*39–41*). Whether IL-2 elicits such signaling programs therefore suppresses osteoclast formation and restores bone mass *in vivo* remains unknown.

Here, we investigated the therapeutic potential and mechanistic basis of therapeutic low-dose IL-2 in reversing ovariectomy (OVX)-induced bone loss. We found that IL-2 functioned as a central regulator of skeletal-immune homeostasis by acting on three coordinated cellular compartments in mice and validated them in human postmenopausal patient samples. We found that IL-2 promoted a CXCR3⁺ T_REG_ population in bone marrow in OVX-mice. IL-2 treatment also reprogrammed bone marrow ILC2s into a regulatory IL-10 and calcitonin gene-related peptide (CGRP) producing state required for skeletal protection. Finally, we found that IL-2 directly engaged osteoclast precursors to activate a STAT1-IRF1 interferon program that suppressed osteoclastogenesis. Collectively, these findings define a multilayered IL-2-responsive network that integrates adaptive immunity, innate lymphoid cell rewiring, and direct myeloid inhibition to restore bone mass. Our data suggest that repurposing low-dose IL-2 may offer a viable immunotherapy to reestablish skeletal integrity in postmenopausal osteoporosis.

## RESULTS

### Therapeutic interleukin-2 protects mice from OVX-induced bone loss

To determine whether immune modulation can restore bone integrity after postmenopausal osteoporosis is established, we used an ovariectomy (OVX) model. Therapeutic low-dose IL-2 (30,000 I.U.) (*42, 43*) was administrated four weeks after surgery and continued for additional four weeks (**Fig. 1A**). High-resolution microcomputed tomography (µCT) imaging of distal femurs showed that IL-2 treatment significantly restored trabecular structure in OVX-mice compared to untreated controls (**Fig. 1B**). Quantitative morphometry confirmed significant gains in bone mineral density (BMD), bone volume fraction (BV/TV), trabecular thickness (Tb.Th), and trabecular number (Tb.N) relative to untreated OVX-mice (p<0.01) (**Fig. 1C**). Histologic analysis further confirmed the improved bone architecture and reduced tartrate-resistant acid phosphatase-positive (TRAP⁺) osteoclasts in IL-2-treated OVX-mice (**Fig. 1D**). These results suggested that therapeutic low-dose IL-2 treatment significantly reversed postmenopausal bone loss in mice.

**Fig. 1.**
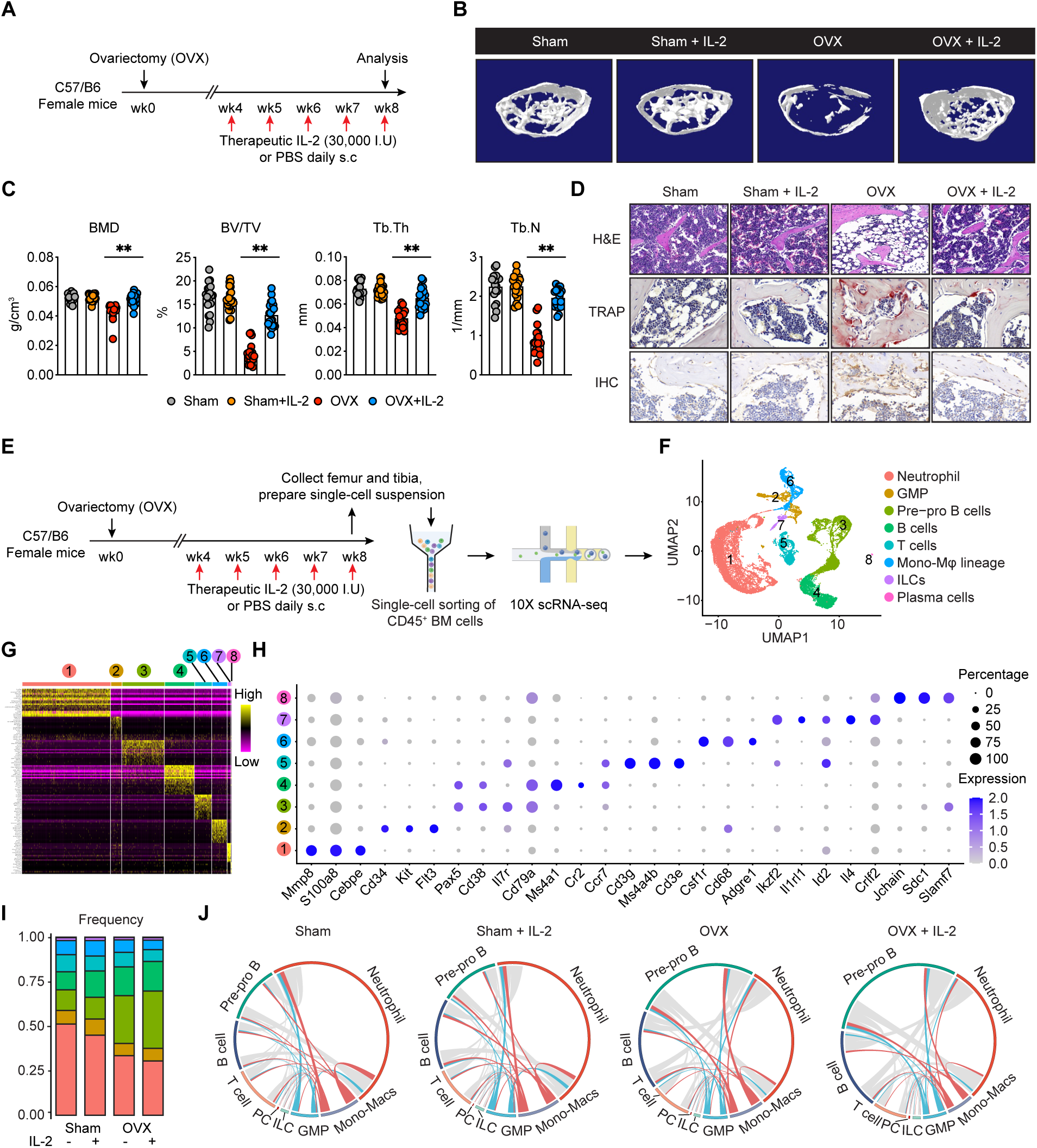
Therapeutic interleukin-2 protects mice from OVX-induced bone loss. **(A)** Schematics of the experiment. WT female mice received daily subcutaneous injections of low-dose IL-2 (30,000 I.U.) or PBS for 4 weeks, 4 weeks after ovariectomy or sham operation. **(B)** Representative 3D μCT images of the distal femurs. **(C)** Quantification of μCT images by BV/TV (bone volume per tissue volume), Tb.N (trabecular number), Tb.Sp (trabecular separation), and Tb.Th (trabecular thickness), n=19-22 mice per group, bar indicates mean, one-way ANOVA, ∗P<0.05, ∗∗P<0.01. **(D)** Representative histological images of bone sections stained with H&E (top), TRAP (middle), and anti-TRAP immunohistochemistry (bottom) to evaluate the abundance of osteoclasts on bone surface. **(E)** Schematics of the single-cell RNA sequencing experiment. CD45^+^ cells were sorted from the bone marrow of the indicated groups for 10x scRNA-seq analysis. **(F)** UMAP visualization of 8 major cell clusters identified in the CD45^+^ bone marrow cells. **(G-H)** Heatmap **(G)** and dot plot **(H)** showing the expression of lineage-defining marker genes used to annotate the clusters. **(I)** Stacked bar graph showing the frequency distribution of cell subsets between groups. **(J)** Circos plots showing the predicted cell-cell interaction networks and interaction strengths among immune clusters in each condition. Interactions unique to monocyte-macrophage lineage cells are shown in red, interactions unique to GMP cells are shown in blue and other interactions are shown in gray.

Given that osteoclasts are derived from the hematopoietic lineage and regulated by local immune signals, we hypothesized that IL-2 exerts its effects by remodeling the bone marrow immune dynamics. To characterize IL–2-induced alterations, we performed single–cell RNA sequencing (scRNA–seq) on CD45⁺ cells isolated from the bone marrow of sham and OVX-mice treated with or without IL-2 (**Fig. 1E, fig. S1**). We recovered 23,368 cells for downstream unsupervised clustering analysis (**Fig. 1F**). Unsupervised clustering of the integrated dataset identified eight distinct immune cell populations, including neutrophils, T cells, B cells, Monocyte-Macrophage lineages and innate lymphoid cells (**Fig. 1F-H**). To examine the remodeling of bone marrow immune landscape by IL-2 administration, we analyzed the global transcriptomic shifts across conditions. Overall composition of CD45^+^ immune cells and their intercellular crosstalk in the marrow changed markedly between groups (**Fig. 1I and J**). This suggests that low-dose therapeutic IL-2 substantially changed the bone marrow immune dynamics.

### IL-2 drives bone-resident CXCR3^+^ regulatory T cells that inhibit osteoclastogenesis

We next investigated the specific T cell subsets responsible for IL-2-mediated bone protection. A focused scRNA-seq analysis of cluster-5 T lymphocytes revealed a significant enrichment of regulatory T cells (T_REG_) with high expression of *Il2ra*, *Ctla4*, *Foxp3* and *Ikzf2* in CD4^+^ T cells in the mouse bone marrow (**Fig. 2A and fig. S2A**). This is consistent with previous reports that as many as 25% of all CD4^+^ T cells in normal human and mouse bone marrow are CD25^+^ T_REG_ cells that help maintain bone homeostasis (*44–46*). As expected, therapeutic IL-2 treatment drastically increased the abundance of bone marrow T_REG_ (BM-T_REG_) in OVX-mice (**Fig. 2B**). Flow cytometry further confirmed more than a two-fold increase in T_REG_ frequency in CD4^+^ T cells after IL-2 administration in OVX-mice compared to the untreated controls (**Fig. 2C and fig. S2B**).

**Fig. 2.**
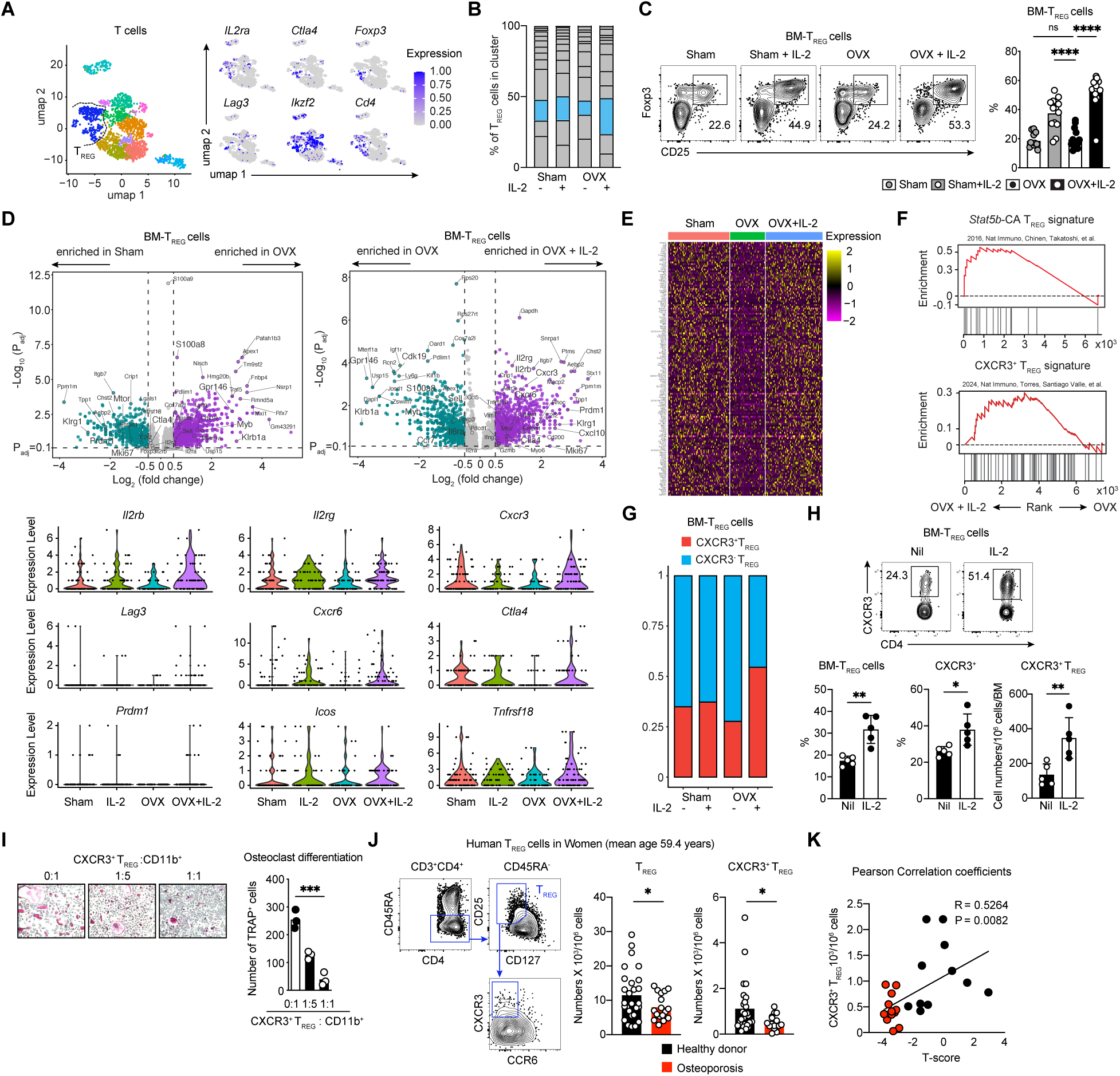
IL-2 drives bone resident CXCR3^+^ regulatory T cells that inhibit osteoclastogenesis. **(A)** UMAP visualization of bone marrow T cell subsets. Feature plots showing the expression of canonical T_REG_ markers *Il2ra, Ctla4, Foxp3, Lag3, Ikzf2,* and *Cd4.* **(B)** Frequency distribution of T cell sub-clusters across Sham and OVX mice treated with vehicle or IL-2, blue indicates T_REG_ cells. **(C)** Representative flow cytometry plots (left) and quantification (right) of CD25^+^Foxp3^+^ T_REG_ frequencies in the BM of Sham, Sham + IL-2, OVX, and OVX + IL-2 treated mice (n=13-15 per group). **(D)** Upper panel, volcano plots showing differentially expressed genes (DEGs) in BM-T_REG_ between Sham and OVX (left) or OVX and OVX + IL-2 (right). Bottom panel, violin plots from scRNA-seq data showing the expression of genes in the indicated group. **(E)** Heatmap of DEGs across the indicated groups. **(F)** Gene Set Enrichment Analysis (GSEA) showing enrichment of *Stat5b*-CA signatures and CXCR3*^+^* T_REG_ signatures in the OVX + IL-2 group compared to OVX control. **(G)** Stacked bar graph showing the ratio of CXCR3^+^ (Red) versus CXCR3^-^ T_REG_ (Blue) from scRNA-seq data. **(H)** WT mice received daily subcutaneous injections of low-dose IL-2 (30,000 I.U.) or PBS for 4 weeks until further analysis. Representative flow cytometry plots (upper) and frequencies and numbers of total BM-T_REG_ and CXCR3^+^ T_REG_ and (bottom), n=5 per group, data are means ± SD, t-test, ∗P<0.05, ∗∗P<0.01. **(I)** CXCR3^+^CD25^+^CD4^+^ T_REG_ cells and bone marrow CD3^-^B220^-^NK1.1^-^CD11b^+^ cells were co-cultured as indicated ratios in the presence of 20 ng/ml M-CSF and 50 ng/ml RANKL for 6 days. Osteoclast formation was assessed by TRAP staining. Representative images (left) and quantification (right) of TRAP^+^ multinucleated (nuclei>3) cells. Representative of at least 2 independent experiments. One-way ANOVA, ∗∗∗P<0.001. **(J)** Flow cytometry analysis of peripheral blood T_REG_ (CD3^+^CD4^+^CD25^+^CD127^low^) and CXCR3^+^ T_REG_ in healthy donors (n=27) compared to patients with osteoporosis (n=17). T-test, data are means ± SD. ∗P<0.05. **(K)** Pearson correlation analysis between CXCR3^+^ T_REG_ and clinical samples with available BMD T-score, n=24, black indicates healthy donor, red indicates osteoporosis patients.

To examine T_REG_ cells and determine how therapeutic IL-2 may influence their transcription and function in the bone microenvironment in the context of OVX-induced bone loss, we performed differential gene expression profiling of BM-T_REG_ cells (**Fig. 2D**). Transcriptomic analysis revealed that IL-2 treatment significantly reversed the global expression pattern of BM-T_REG_ cells (**Fig. 2E**). Genes including *S100a8*, *Gpr146*, *Apex1*, *Klrb1a* and *Myb* that were enriched in BM-T_REG_ from OVX mice were significantly downregulated in BM-T_REG_ cells isolated from IL-2 treated OVX-mice. By contrast, *Il2rb*, *Il2rg*, *Cxcr3*, *Cxcr6*, *Prdm1*, *Klrg1*, *Mki67* and *Ctla4* were among the genes that were highly upregulated in BM-T_REG_ cells from IL-2 treated relative to untreated OVX-mice (**Fig. 2D-E**). When aligning our scRNA-seq data to the published dataset by gene-set enrichment analysis, we found significant induction of a *Stat5b*-activated signature in BM-T_REG_ cells, confirming the increased IL-2 receptor signaling from IL-2 treated OVX-mice (**Fig. 2F**) (*47*). Moreover, we observed that BM-T_REG_ cells from IL-2 treated OVX-mice showed markedly enriched programs characteristic of CXCR3⁺ effector T_REG_ cells (**Fig. 2F**), a population that were associated with tissue protection in inflammatory settings (*48*). Notably, these CXCR3⁺ T_REG_ cells were abundantly enriched in the BM microenvironment after IL-2 administration (p<0.05) (**Fig. 2G-H**).

To test whether CXCR3^+^ T_REG_ cells directly influence osteoclast differentiation, we cocultured flow cytometric sort-purified bone marrow CXCR3^+^ CD25^+^ CD4^+^ T cells with CD11b⁺ monocytes undergoing RANKL-induced osteoclastogenesis wherein CXCR3^+^ T_REG_ cells strongly suppressed osteoclast formation (**Fig. 2I, fig. S2C**). Consistent with the findings in mice, we observed substantial reduction in total T_REG_ and CXCR3⁺ T_REG_ populations in peripheral blood samples collected from postmenopausal osteoporosis patients (OP) compared to samples from age-matched women healthy donors (HD) (mean age OP 60.7 years, HD 58.3 years, p=0.13; total T_REG,_ p<0.05; CXCR3⁺ T_REG_, p<0.05) (**Fig. 2J, fig. S2D**). In addition, correlation analysis revealed a significant and positive association between the bone density T-score and CXCR3^+^ T_REG_ cells in our cohort (p=0.0082, r=0.5264) (**Fig. 2K**). Together, these results indicate that therapeutic IL-2 treatment expands a specialized CXCR3⁺ T_REG_ population to inhibit osteoclastogenesis.

### *Il10* and *Calca*-expressing ILC2 cells restrain osteoclastogenesis in response to IL-2

To determine additional cellular mechanisms independent of adaptive immunity that underscore the significantly improved bone mass, we treated *Rag1*^-/-^ mice, wherein T and B cells were deficient (*49*), with the same therapeutic IL-2 regimen (**Fig. 3A**). µCT imaging analysis of distal femurs showed that daily IL-2 administration significantly ameliorated bone loss in OVX-*Rag*^-/-^mice, which were quantified by increased BMD, bone volume fraction, trabecular thickness and number (Tb.N) of treated and untreated mice (p<0.01) (**Fig. 3B-C**). Improved bone integrity was associated with a reduced number of TRAP^+^ osteoclasts in metaphysis region of distal femur (**Fig. 3D**). To examine the cellular mechanisms, we analyzed our scRNA-seq dataset and identified a population of innate lymphoid cells (ILCs) that were accumulated in the bone marrow of *Rag1*^-/-^mice (**Fig. 3E, fig. S3A**). scRNA-seq analysis further revealed that IL-2 treatment drove a specific expansion of type 2 ILCs (ILC2s), characterized by high expression of *Gata3*, *Il1rl1* (ST2), *Ikzf2, Id2, IL4, Il2ra and Ramp1* in mice received OVX surgery (p=0.021, **Fig. 3E-G**). We further verified the expansion of ILC2s by IL-2 treatment using flow cytometry (**Fig. 3H, fig. S3B**) and found a significant increase of BM-ILC2 cells in both wild-type (p<0.001) and *Rag1*^-/-^ deficient mice (p<0.0001) compared to those OVX-mice that did not receive treatment (**Fig. 3I**).

**Fig. 3.**
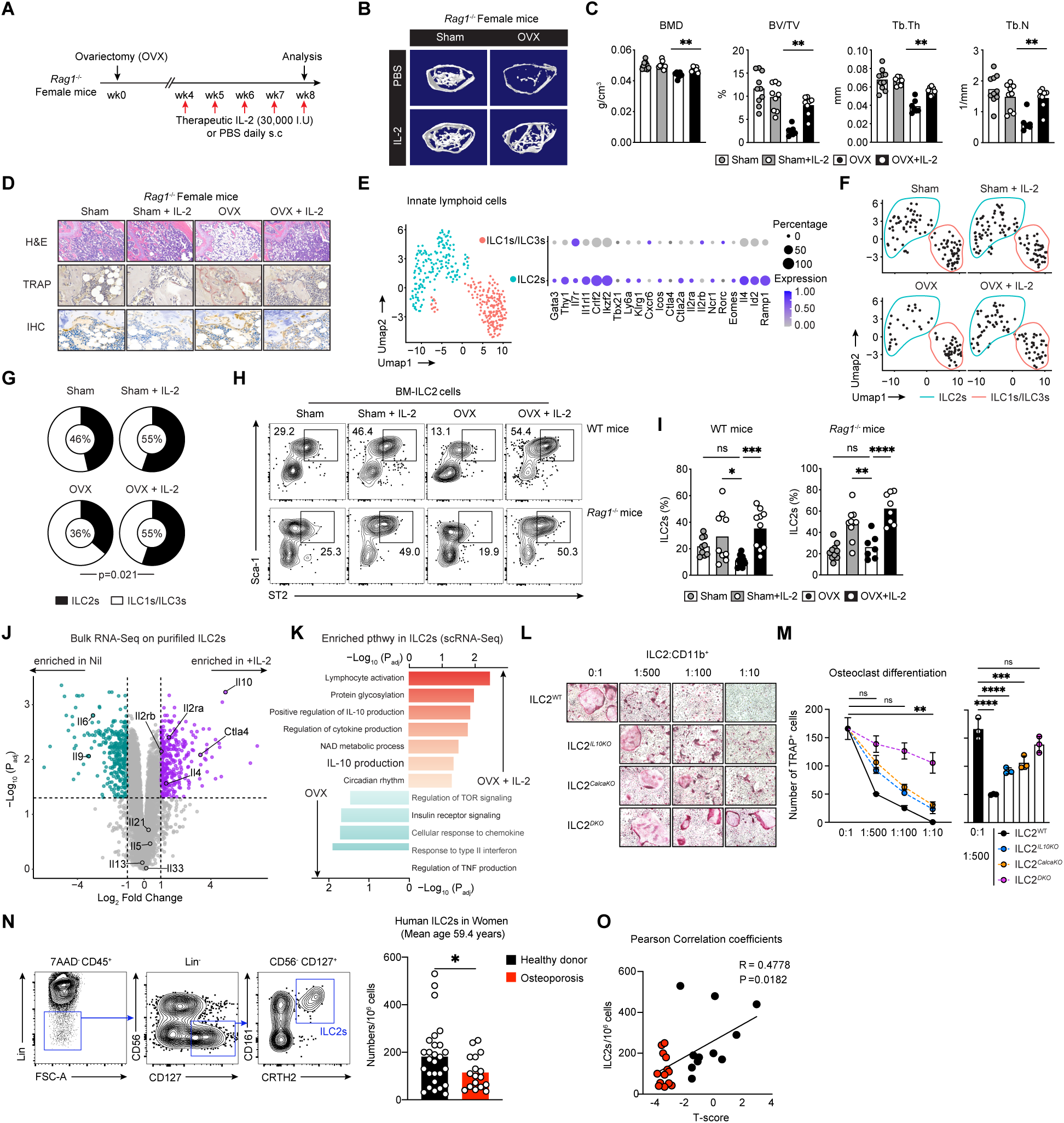
*Il10* and *Calca*-expressing ILC2 cells restrain osteoclastogenesis in response to IL-2. **(A)** Schematics of the experiment. *Rag1^-/-^* female mice received daily subcutaneous injections of low-dose IL-2 (30,000 I.U.) or PBS for 4 weeks, 4 weeks after ovariectomy or sham operation. **(B)** Representative 3D μCT images of the distal femurs. **(C)** Quantification of μCT images by BV/TV (bone volume per tissue volume), Tb.N (trabecular number), Tb.Sp (trabecular separation), and Tb.Th (trabecular thickness), n=7-10 per group, bar indicates mean, one-way ANOVA, ∗P<0.05, ∗∗P<0.01. **(D)** Representative histological images of bone sections stained with H&E (top), TRAP (middle), and anti-TRAP immunohistochemistry (bottom) to evaluate the abundance of osteoclasts on bone surface. **(E)** UMAP visualization of bone marrow innate lymphoid cell (ILC) subsets from scRNA-seq data (Left) with key genes identifying ILC2s and ILC1s/ILC3s subclusters (Right). **(F)** Split UMAP visualization showing the distribution of bone marrow ILCs in each treatment group. **(G)** Pie charts showing the ratio of ILC2s to ILC1s/ILC3s. **(H-I)** Representative flow cytometry plots **(H)** and the quantification of BM-ILC2s (Lin^-^CD45^+^CD127^+^ST2^+^Sca-1^+^) in WT and *Rag1^-/-^* mice **(I)**. **(J)** Volcano plot of bulk RNA-seq data from purified ILC2s showing differential expression between control and IL-2 treated groups. **(K)** GO Pathway enrichment analysis of ILC2s comparing OVX vs. OVX + IL-2 groups from scRNA-seq data. (L-M) FACS sorted bone marrow CD3^-^B220^-^NK1.1^-^CD11b^+^ cells and ILC2s (Lin^-^CD45^+^CD127^+^Sca-1^+^ST2^+^) from WT, *Il10^-/-^, Calca^-/-^* and *Il10^-/-^Calca ^-/-^* (DKO) were cocultured at indicated ratios, in the presence of 20 ng/ml M-CSF and 50 ng/ml RANKL. Osteoclast formation was evaluated by TRAP staining. Representative images **(L)** and quantification **(M)** of TRAP^+^ multinucleated (nuclei > 3) cells. Representative of at least 3 independent experiments. Bar indicates mean, one-way ANOVA, ∗∗P<0.01, ∗∗∗P<0.001, ∗∗∗∗P<0.0001. **(N)** Flow cytometry analysis of peripheral blood ILC2s in healthy donors (n=27) compared to patients with osteoporosis (n=17). T-test, ∗P<0.05. **(O)** Pearson correlation analysis between ILC2s and clinical samples with available BMD T-score, n=24, black indicates healthy donor, red indicates osteoporosis patients.

Bulk RNA-seq analysis of *ex vivo* purified ILC2 cells revealed that IL-2 treatment only modestly increased *Il4* expression with negligible or negative effects on other canonical type 2 cytokine genes including *Il5*, *Il9*, and *Il13* (**Fig. 3J**). By contrast, IL-2 markedly upregulated regulatory-like genes such as *Il10* and *Ctla4* in ILC2s (**Fig. 3J**). Consistently, pathway analysis of scRNA-seq data confirmed that therapeutic IL-2 reprogramed ILC2s in OVX-mice toward an IL-10-producing state *in vivo* (**Fig. 3K**). Enzyme-linked immunosorbent assay (ELISA) of culture supernatants from *ex vivo* IL-2 treated ILC2s further confirmed their robust production of IL-10 (**fig. S3C**).

To determine whether IL-10-producing ILC2s regulate osteoclastogenesis, we collected cell culture supernatants with or without IL-10 neutralizing antibody from *ex vivo* expanded ILC2s and added into purified CD11b^+^ monocytes under OC formation for 6 days (**fig. S3D**). As expected, 1% dilution of ILC2s supernatant profoundly inhibited the RANKL-induced OC formation (**fig. S3E**), while this inhibition by supernatant was markedly impaired when IL-10 neutralizing antibody was added into the ILC2 culture (**fig. S3E-F**). Next, we isolated ILC2s from *Il10*^KO^ mice (ILC2*^Il10KO^*) and co-cultured with bone marrow purified CD11b^+^ monocytes at a ratio of 1:500, 1:100 or 1:10, wherein CD11b^+^ cells were differentiated into osteoclasts by M-CSF and RANKL. As expected, wild-type ILC2s (ILC2^WT^) markedly inhibited osteoclast differentiation at a ratio of 1:500, while ILC2*^Il10KO^* cells exhibited largely impaired suppression of osteoclastogenesis from CD11b^+^ cells (**Fig. 3L-M**).

Growing evidence suggest that neuropeptides shape bone remodeling (*50–52*), and ILC2s are known producers of calcitonin gene-related peptide (CGRP) through *Calca* expression (*53–55*), we next tested whether a neuropeptidergic effector contributes to their anti-osteoclastogenic activity. ELISA of culture supernatants from *ex vivo* IL-2 treated ILC2s revealed robust production of CGRP (**fig. S3G**). Moreover, CGRP blockade significantly impaired the magnitude of ILC2 supernatants to inhibit osteoclastogenesis (**fig. S3E-F**). Consistently, ILC2s isolated from *Calca*^KO^ mice (ILC2*^CalcaKO^*) exhibited reduced suppression of osteoclastogenesis in co-culture, resulting in increased osteoclast numbers compared with ILC2^WT^ cells (**Fig. 3L-M**). To examine whether IL-10 and CGRP act cooperatively, we generated *Il10*^-/-^ *Calca*^-/-^ double knockout mice and isolated ILC2s (ILC2*^DKO^*) for co-culture under RANKL induced osteoclastogenesis. Notably, ILC2*^DKO^* cells exhibited a profound loss of inhibitory function, failing to suppress osteoclast differentiation even at higher ILC2-to-CD11b^+^ cell ratios (1:100 or 1:10) (**Fig. 3L-M**). These results indicate that IL-10 and CGRP operate cooperatively and are both required for ILC2-mediated protection. In our postmenopausal osteoporosis patient cohorts, we observed a significant decrease of ILC2s in patients compared to age-matched healthy donors (p<0.05) (**Fig. 3N**), and a positive correlation between ILC2s and T-score of the bone density (p=0.0182, r=0.4778) (**Fig. 3O**). Together, these results highlight critical and previously underappreciated roles for IL-10- and CGRP-producing ILC2s in protecting against bone loss, effects that can be further potentiated by IL-2 treatment.

### IL-2 directly inhibits osteoclastogenesis

To determine whether IL-2-mediated protection against bone loss extends beyond lymphocyte-dependent mechanisms, we depleted ILCs in OVX *Rag1*^-/-^ mice using anti-CD90 antibody (*56*) (**Fig. 4A**). In this system, daily therapeutic IL-2 was administered for four weeks after establishment of OVX-induced bone loss (**Fig. 4A**). Sufficient depletion of ILC populations in *Rag1*^-/-^ mice was confirmed (**fig. S4A**). Interestingly, therapeutic IL-2 treatment ameliorated OVX induced-bone loss in the absence of T cells and ILCs, although the magnitude of protection was smaller compared to the T and ILC sufficient condition (**Fig. 4B-D**). These results suggested that IL-2 may directly inhibit osteoclastogenesis within the myeloid lineage.

**Fig. 4.**
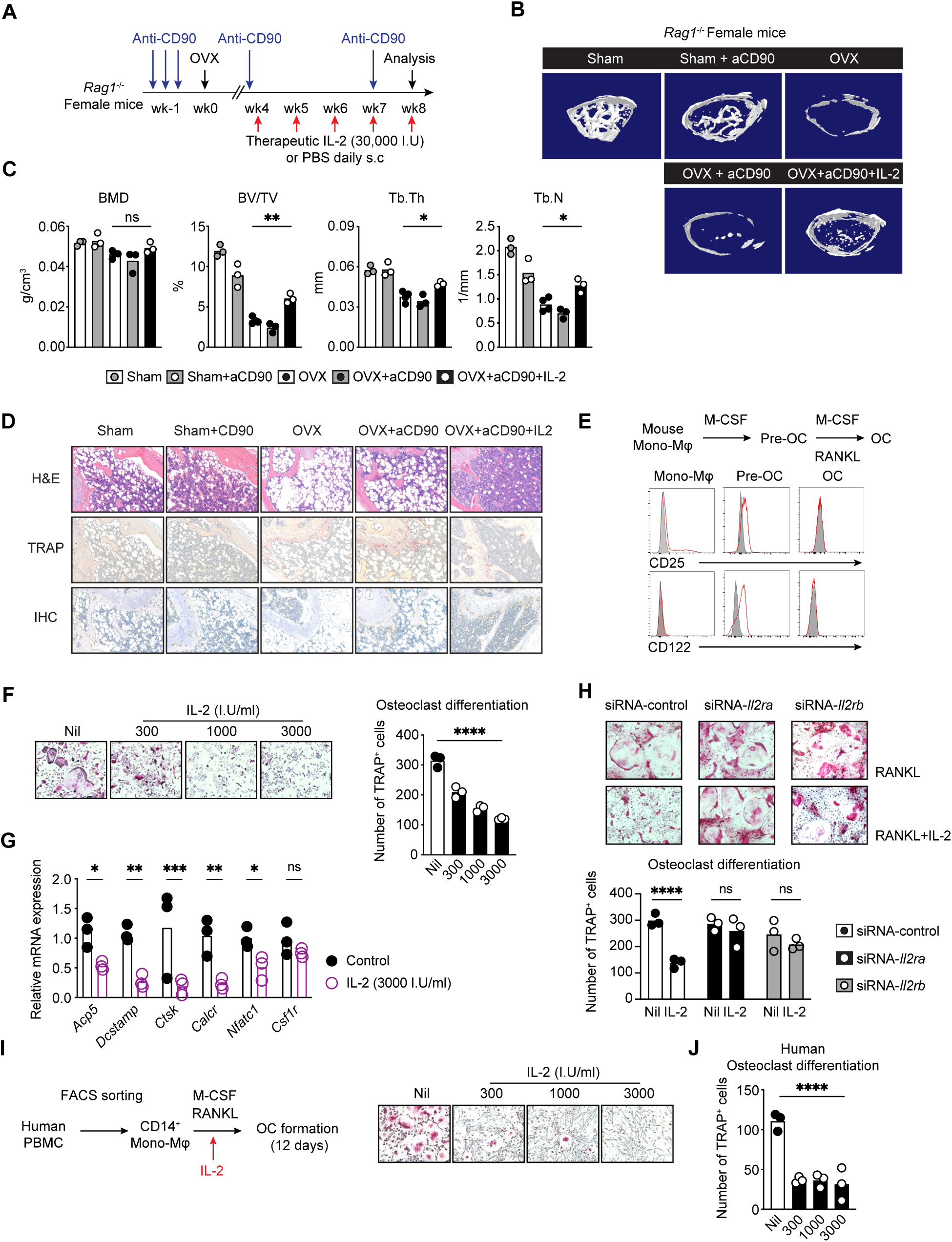
IL-2 directly inhibits osteoclastogenesis. **(A)** Schematics of the experiment to deplete ILCs using anti-CD90 antibody in *Rag1^-/-^* mice with OVX or Sham operation. **(B)** Representative 3D μCT images of the distal femurs. **(C)** Quantification of μCT images by BV/TV (bone volume per tissue volume), Tb.N (trabecular number), Tb.Sp (trabecular separation), and Tb.Th (trabecular thickness), n=3-4 per group, bar indicates mean, one-way ANOVA, ∗P<0.05, ∗∗P<0.01. **(D)** Representative histological images of bone sections stained with H&E (top), TRAP (middle), and anti-TRAP immunohistochemistry (bottom) to evaluate the abundance of osteoclasts on bone surface. **(E)** FACS histogram of CD25 and CD122 expressions during osteoclastogenesis. Grey, isotype control. Red, CD25 or CD122 expression. **(F)** Differentiation of bone marrow CD3^-^B220^-^NK1.1^-^CD11b^+^ cells into osteoclasts in the presence of RANKL and M-CSF, with indicated concentrations of IL-2 added at the start of cell culture. Osteoclast formation was evaluated by TRAP staining. Representative images (left) and quantification (right) of TRAP^+^ multinucleated (nuclei > 3) cells. Representative of at least 3 independent experiments. One-way ANOVA, ∗∗∗∗P<0.0001. **(G)** qPCR of osteoclast-related gene expression in CD3^-^B220^-^NK1.1^-^CD11b^+^ cells 4 days after stimulation with RANKL and M-CSF. Black dots, control, red dots, IL-2 treatment (3,000 I.U/ml). Representative of at least 3 independent experiments. Bar indicates mean. T-test, ∗P<0.05, ∗∗P<0.01, ∗∗∗P<0.001, ∗∗∗∗P<0.0001. **(H)** Cultured CD3^-^B220^-^NK1.1^-^CD11b^+^ cells were transfected with siRNA targeting *Il2ra* or *Il2rb* following RANKL-induced osteoclastogenesis with or without IL-2 treatment (3000 I.U/ml). Representative TRAP images (top) and quantification of TRAP^+^ multinucleated cells (nuclei > 3, bottom). Representative of at least 3 independent experiments. Bar indicates mean. T-test, ∗∗∗∗P<0.0001. **(I-J)** *In vitro* differentiation of human osteoclasts from PBMCs. FACS sorted CD14^+^ monocytes were differentiated into osteoclasts with recombinant human IL-2. Representative TRAP images **(I)** and quantification **(J)**. Representative of at least 3 independent experiments. Bar indicates mean. One-way ANOVA, ∗∗∗∗P<0.0001.

To test this hypothesis, we first examined IL-2 receptor expression during RANKL-induced osteoclastogenesis and found that both IL-2 receptor α (CD25) and β (CD122) were strongly upregulated in osteoclast progenitors (pre-OCs) three days after induction but declined as cells matured into terminal osteoclasts (**Fig. 4E**). Next, we differentiated FACS-sorted BM B220^-^CD3^-^NK1.1^-^CD11b^+^ Mono-Mφ by M-CSF and RANKL (**fig. S4B**) and found that titrating IL-2 at the beginning of cell culture resulted in a dose-dependent blockade of osteoclast formation (**Fig. 4F**), an effect further supported by reduced bone resorption from pit formation assays (**fig. S4C**). Correspondingly, IL-2 treatment strongly suppressed the expression of key osteoclastogenic genes, including *Acp5*, *Dcstamp*, *Ctsk*, *Calcr*, and *Nfatc1* (p<0.05) (**Fig. 4G**).

To confirm the direct regulation is through IL-2 receptor signaling, we performed siRNA knockdown of *Il2ra* or *Il2rb* on Mono-Mφ undergoing RANKL-induced osteoclastogenesis and observed significantly impaired inhibition of osteoclast differentiation and gene expression by IL-2 compared to the controls with intact IL-2 receptors (**Fig. 4H, fig. S4D-E**). These results indicate that IL-2 directly regulates osteoclastogenesis. Finally, in cultures of FACS-sorted human CD14⁺ Mono-Mφ from PBMCs (**fig. S4F**), IL-2 induced a significant, dose-dependent reduction in osteoclast formation over a 12-day differentiation period (p<0.0001) (**Fig. 4I-J**). Together, these studies establish IL-2 as a direct negative regulator of osteoclastogenesis in both mice and humans.

### IL-2 reprograms RANKL-stimulated monocytes

To examine IL-2-induced changes in gene expression during osteoclastogenesis, we performed high-throughput RNA sequencing (RNA-seq) on flow cytometric sort-purified B220^-^CD3^-^NK1.1^-^CD11b^+^ Mono-Mφ undergoing RANKL-induced differentiation, with or without IL-2 treatment (**Fig. 5A, fig. S5**). Principal component analysis (PCA) revealed a clear segregation of IL-2-treated RANKL-stimulated cells from those stimulated with RANKL alone (**Fig. 5B**). Pathway analysis demonstrated that IL-2 markedly downregulated osteoclastogenic transcriptional programs (**Fig. 5C**). This downregulation includes pathways required for osteoclast differentiation, bone remodeling, bone resorption, calcium signaling, response to reactive oxygen species (ROS) and cell-cell fusion (**Fig. 5C**). Moreover, IL-2 treatment directly upregulated genes involved in response to interferon beta, cytokine mediated signaling, type I interferon production, mononuclear cell proliferation and interferon mediated signaling (**Fig. 5C**). Specifically, four days after RANKL stimulation, canonical osteoclast-associated genes including *Ctsk*, *Mmp9*, *Ocstamp*, *Dcstamp*, and *Acp5* were highly upregulated relative to unstimulated controls as expected (**Fig. 5D**). By contrast, IL-2 treatment suppressed key genes of the osteoclast fusion and resorption machinery, such as *Dcstamp*, *Ocstamp*, *Ctsk* and *Acp5* indicating a profound IL-2-mediated transcriptional reprogramming that opposes osteoclastogenesis (**Fig. 5C-D**). This inhibitory effect was further illustrated by RNA-seq based heatmap visualization incorporating all experimental conditions (**Fig. 5E**), as well as the decreased counts-per-million (CPM) expression of osteoclast signature genes shown in **Fig. 5F**.

**Fig. 5.**
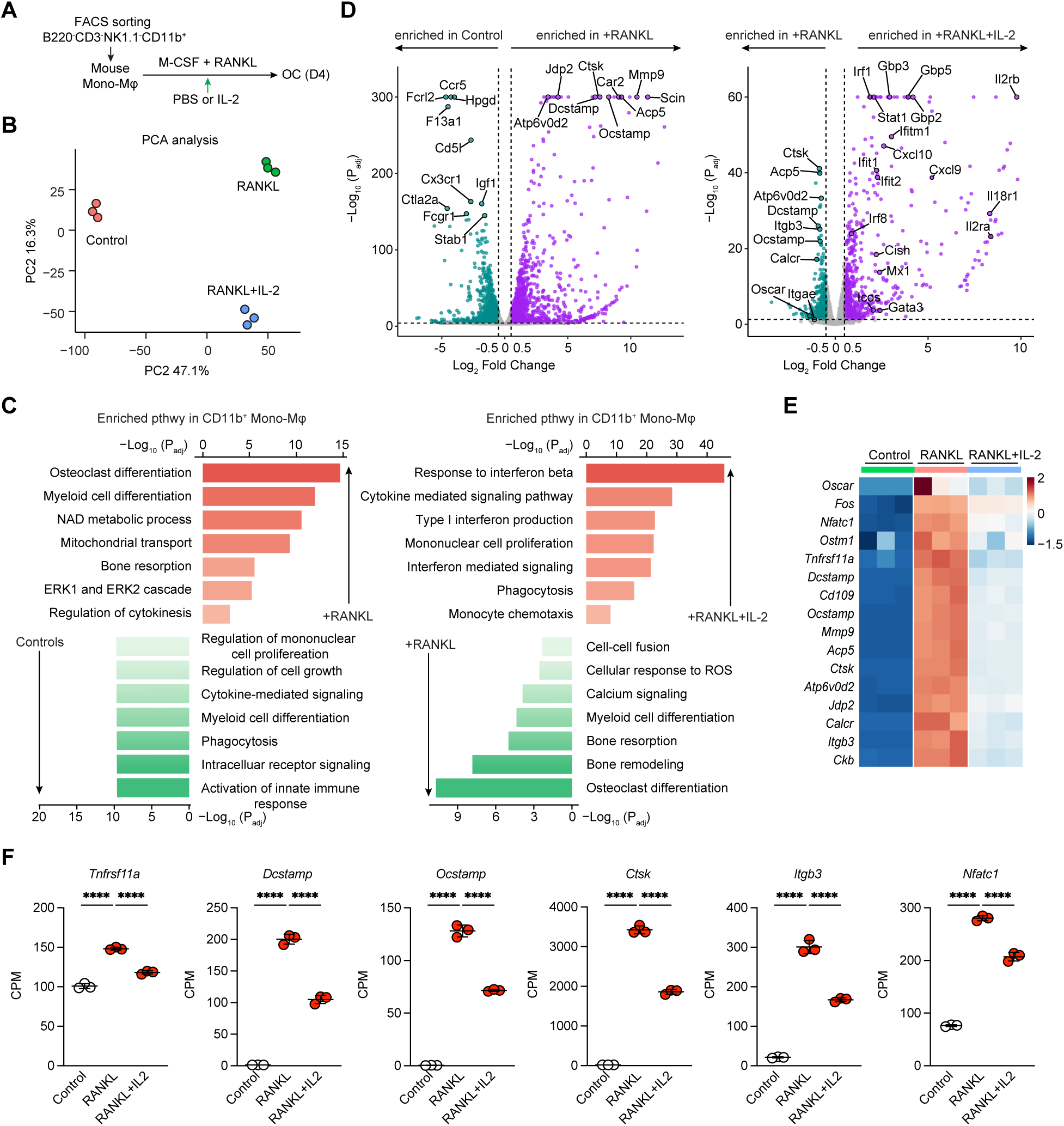
IL-2 reprograms RANKL-stimulated monocytes. **(A)** Schematics of the bulk RNA-seq experiment. Mouse bone marrow CD3^-^B220^-^NK1.1^-^CD11b^+^ cells were FACS sort-purified and differentiated into osteoclast in the presence of with RANKL (50 ng/ml) and M-CSF (20 ng/ml) with or without IL-2 (3,000 I.U/ml) treatment for 4 days (n=3 biological replicates per group). **(B)** PCA analysis of control, RANKL, and RANKL + IL-2 groups. **(C)** GO pathway enrichment analysis of cells between control and RANKL (left), or RANKL and RANKL+IL-2 groups (right). **(D)** Volcano plots of RNA-seq from different groups showing differential gene expression between control and RANKL, or RANKL and RANKL+IL-2 groups. **(E)** Heatmap of canonical osteoclast marker genes including *Nfatc1, Ctsk, Dcstamp, Acp5* among all three groups **(F)** Normalized expression (CPM) of key osteoclastogenic genes (*Tnfrsf11a, Dcstamp, Ocstamp, Ctsk, Itgb3, Nfatc1*) in the indicated groups. Data are means ± SD. One-way ANOVA, ∗∗∗∗P<0.0001.

### Activation of IFN-program is essential for the IL-2-mediated intrinsic suppression of osteoclastogenesis

PCA analysis indicated that IL-2 redirected differentiating monocytes toward a transcriptional state distinct from both undifferentiated controls and mature osteoclasts (**Fig. 5B**). To define the transcriptional programs underlying this IL-2-mediated diversion, we performed pathway analysis on differentially expressed genes. A prominent theme that emerged was the repeated enrichment of interferon-related pathways and interferon-stimulated genes by IL-2 treatment (**Fig. 6A-E, fig. S6A**). Within this program, *Irf1* and *Stat1* exhibited a 2-3-fold increase in IL-2-treated cells compared with RANKL-stimulated controls, whereas expression of *Ifnar1* remained unchanged between the two conditions (**Fig. 6F**).

**Fig. 6.**
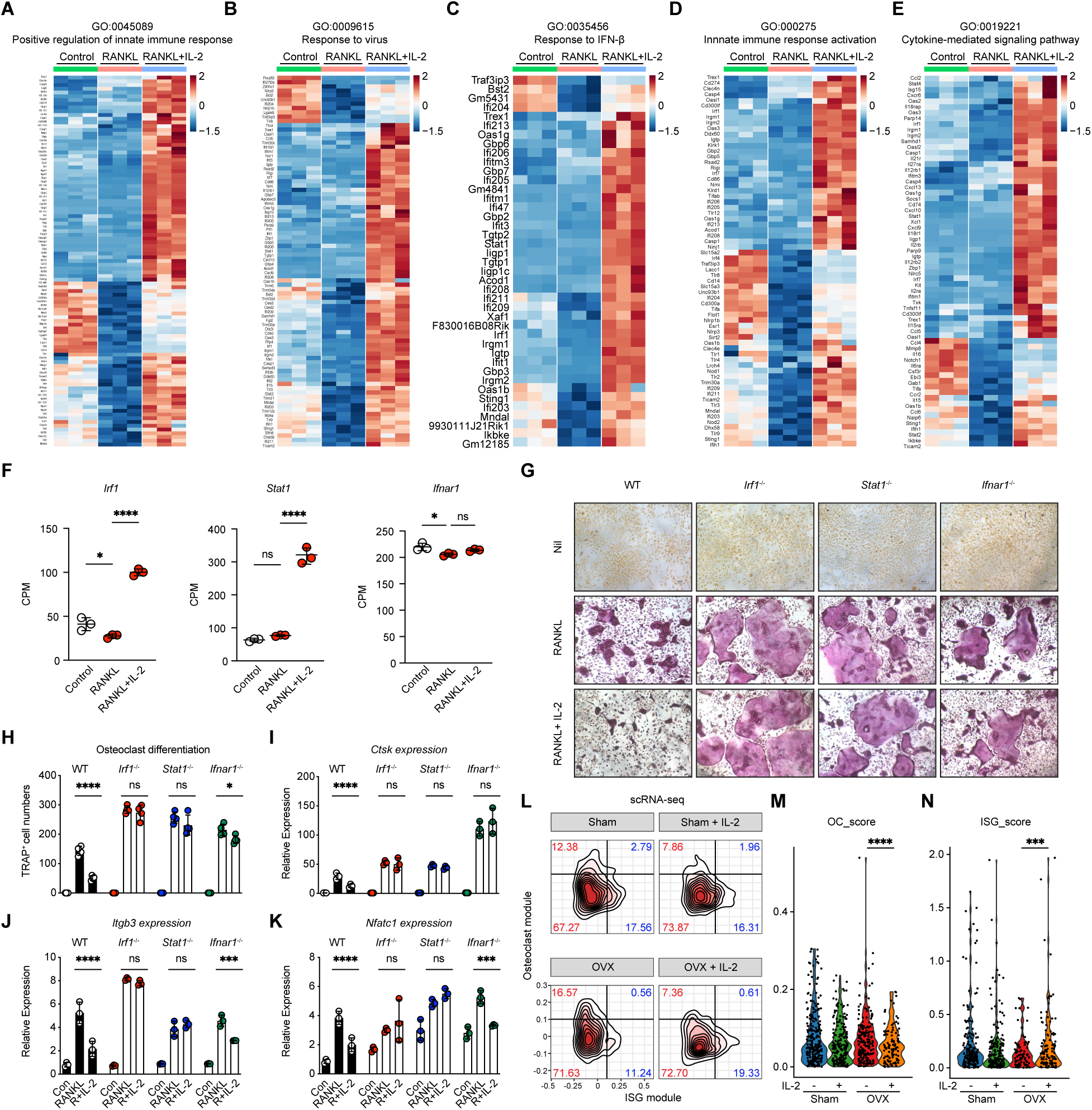
Activation of IFN-program is essential for the IL-2-mediated intrinsic suppression of osteoclastogenesis. **(A-E)** Heatmaps of multiple GO pathways from bulk RNA-seq data. RANKL+IL-2 treated group shows enrichment in multiple GO pathways: positive regulation of innate immune response **(A)**, Response to virus **(B)**, Response to IFN-β **(C)**, Innate immune response activation **(D)**, and Cytokine-mediated signaling pathways **(E)**. **(F)** Normalized expression (CPM) of interferon-related transcription factors (*Irf1, Stat1*) and receptor (*Ifnar1*) in the indicated groups. One-way ANOVA, ∗P<0.05, ∗∗∗∗P<0.0001. **(G-K)** CD3^-^B220^-^NK1.1^-^CD11b^+^ cells were sorted from WT, *Irf1^-/-^*, *Stat1^-/-^,* and *Ifnar1^-/-^* mice and differentiated into osteoclasts with RANKL and M-CSF in the presence or absence of IL-2. **(G)** Representative TRAP staining images. **(H)** Quantification of TRAP^+^ multinucleated cells. Representative of at least 3 independent experiments. **(I-K)** Relative mRNA expression of osteoclast marker genes *Ctsk* **(I)**, *Itgb3* **(J)**, and *Nfatc1* **(K)** assessed by qPCR. **(L)** Contour plot of the OC progenitor-enriched clusters from scRNA-seq dataset showing Osteoclast (OC) module or Interferon-Stimulated Gene (ISG) module. **(M-N)** Violin plots showing the Osteoclast (OC) Score **(M)** and Interferon-Stimulated Gene (ISG) Score **(N)** within the OC progenitor-enriched clusters in Sham and OVX mice treated with or without IL-2. One-way ANOVA, ∗∗∗∗P<0.0001.

We next tested whether the IL-2-mediated direct inhibition of osteoclastogenesis requires *Irf1*-*Stat1* signaling. To address this, we generated osteoclasts from bone marrow cells isolated from *Irf1*^−/−^, *Stat1*^−/−^, and *Ifnar1*^−/−^ mice (**Fig. 6G**). Consistent with previous studies (*39–41*), loss of any of these genes resulted in significantly increased RANKL-induced osteoclast formation compared to cultures with WT cells (**Fig. 6G**). Notably, while IL-2 potently suppressed osteoclastogenesis in WT cells, this inhibitory effect was significantly abolished in *Irf1*^−/−^ and *Stat1*^−/−^ cultures, wherein comparable TRAP^+^ cells were found in IL-2-treated or RANKL-only conditions (**Fig. 6G and 6H**). By contrast, IL-2 continued to potently suppress osteoclastogenesis in *Ifnar1*^−/−^ cells (p<0.05), indicating that IFNAR1 is dispensable for this effect. Furthermore, loss of *Stat1-Irf1* signaling eliminated IL-2’s ability to suppress *Ctsk*, *Itgb3* and *Nfatc1* expression (**Fig. 6I-K**). Unsupervised scRNA-seq analysis of myeloid clusters 1, 2, and 6 (**Fig. 1F**) resolved 12 transcriptionally distinct subpopulations (**fig. S6B-C**). Among these, clusters 3, 10, and 11 contained cells with hallmarks of osteoclast progenitors, characterized by high expression of *Csf1r*, *Rgs10*, *Ccr2*, and *Cx3cr1*, and low expression of *Ly6g*, *S100a8*, *S100a9*, and *Cxcr2* (**fig. S6D-E**). Notably, therapeutic IL-2 markedly reduced the osteoclast scores within these OC progenitor-enriched clusters (**Fig. 6L, fig. S6F**), while increased interferon-stimulated gene (ISG) scores in IL-2-treated OVX mice compared to untreated OVX controls (**Fig. 6M-N**). Collectively, these data demonstrate that therapeutic IL-2 directly regulates osteoclastogenesis through interferon dependent-signaling to intrinsically suppress osteoclast lineage commitment.

## DISCUSSION

Postmenopausal osteoporosis remains a major global health challenge, driven primarily by excessive osteoclast activity and progressive loss of trabecular bone (*1–3*). Recent advances in osteoimmunology revolutionized our understanding of osteoporosis pathophysiology and demonstrated the key role of bone-immune intercellular circuits in influencing the balance between bone formation and resorption (*57, 58*). However, novel immunotherapy that can overcome hurdles from current antiresorptive and anabolic therapies such as long-term side effects (*4*), plateauing efficacy, or rapid bone loss upon discontinuation (*5, 6*), remains poorly developed. Here, we examined a clinically accessible low-dose IL-2 based immunotherapy in mouse bone loss models and found that therapeutic use of IL-2 restored trabecular bone architecture in mice with OVX-induced bone disruption. Mechanistically, we found that IL-2-mediated protection was mainly through the reprogramming of the bone marrow immune circuits to systemically limit osteoclastogenesis.

Low-dose IL-2 therapy has emerged as a safe and effective approach for the treatment of autoimmune and inflammatory diseases (*19, 21*). The therapeutic rationale is to preferentially expand T_REG_ cells and restore T_REG_-mediated immune balance through engagement of the high-affinity IL-2 receptor α chain (CD25), while limiting the activation of cytotoxic lymphocytes (*36*). Consistent with this concept, low-dose IL-2 has shown clinical benefit in systemic diseases including systemic lupus erythematosus (SLE) (*9–14*), vasculitis (*15*), and graft-versus-host disease (GVHD) (*16*). However, how low-dose therapeutic use of IL-2 shapes tissue-specific T_REG_ programs within distinct disease microenvironments, including bone tissue, remains largely unexplored. Our experiments show that IL-2 treatment expands a CXCR3⁺ T_REG_ compartment that is critical in reversing bone loss. This indicates that expression of certain migratory molecules may be essential for the residency of regulatory cells in the marrow and their access to the chemokine-defined bone remodeling niches to restrain osteoclastogenesis. Consistently, low-dose IL-2 therapy was shown to induce CXCR3 expression on T_REG_ cells in SLE patients (*59*). In this setting, CXCR3⁺ T_REG_ co-express CD38 and HLA-DR and display a highly proliferative phenotype, facilitating their recruitment to sites of acute inflammation (*59*). Notably, CXCR3⁺ T_REG_ was associated with enhanced tissue repair capacity and effective control of inflammation (*48*). In addition, CXCR3 signaling was mechanistically linked to the RANKL-induced bone remodeling (*60*), wherein osteoblast progenitors produced *Cxcl9* ortholog *Cxcl9l* that recruited *Cxcr3.2*⁺ osteoclast progenitors to sites of bone resorption. Therefore, in OVX-induced bone loss model, therapeutic IL-2 may preferentially expand and relocate CXCR3⁺ T_REG_ cells to the sites of bone resorption and locally restrain osteoclastogenesis.

ILC2s express high levels of the high-affinity IL-2 receptor α chain (CD25), preparing them to respond to the low concentrations of IL-2 treatment (*61*). Indeed, our experiments show that therapeutic administration of low-dose IL-2 in OVX-mice induces a substantial accumulation of bone marrow ILC2s. Moreover, IL-2 reprograms ILC2s toward an IL-10- and CGRP-producing state. Loss-of-function experiments using *Il10*-deficient, *Calca*-deficient, and double-deficient mice confirmed the essential roles of *Il10* and *Calca* for ILC2s to restrain osteoclastogenesis. Canonical ILC2s producing type 2 cytokines IL-4 and IL-13 were shown to directly interfere osteoclastogenesis (*62, 63*). Our experiments are distinct from these by showing that a clinically relevant cytokine therapy such as low-dose IL-2 can directly expand ILC2 population *in vivo* and promote their regulatory function in protecting bone mass.

Activated monocytes and macrophages express IL-2 receptors (*33–35*). Our experiments indicated that both CD25 and CD122 were transiently upregulated on pre-osteoclasts during RANKL-induced differentiation. IL-2 treatment during this stage suppressed osteoclast maturation and bone-resorptive activity in both mouse and human cell cultures. RNA-seq profiling revealed that IL-2 redirected differentiating osteoclast precursors toward a distinct interferon-active state characterized by activation of *Stat1* and *Irf1* and repression of osteoclastogenic programs. Genetic ablation of *Stat1* or *Irf1* abolished the inhibitory effects of IL-2, whereas loss of *Ifnar1* showed minimal effects, indicating that IL-2 induces an IFN-like transcriptional program independently of type I IFN autocrine loops. These mechanistic data establish a direct, lineage-intrinsic role for IL-2 in driving osteoclast precursors away from terminal differentiation.

In summary, our work establishes IL-2 as a regulator of skeletal-immune crosstalk that operates through coordinated effects on T_REG_, ILC2s, and osteoclast precursors. By integrating adaptive immunity, innate lymphoid pathways, and lineage-intrinsic reprogramming, therapeutic IL-2 effectively halts osteoclastogenesis and reverses established bone loss in a preclinical postmenopausal bone loss model. These findings uncover an unappreciated axis of immune control over bone remodeling and open new avenues for cytokine-based interventions in osteoporosis and related skeletal disorders.

## MATERIALS AND METHODS

### Human samples

All patients and healthy controls provided written informed consent. 17 postmenopausal osteoporosis patients and 27 age-matched healthy women were enrolled in this study, and their blood samples were collected at Qilu Hospital of Shandong University, Jinan, China. Adult BMD was used to define bone health with calculated T-score and Z-score to report the BMD. Patients with ongoing or previous systemic autoimmune diseases or severe infectious diseases were excluded from the study. All healthy individuals had no known history of any significant systemic diseases, including, but not limited to, autoimmune disease, diabetes, allergic disease, kidney or liver disease, or malignancy. Individual characteristics are provided in Tables S1.

### Mice

All mouse lines were maintained on a C57BL/6J background in SPF (specific pathogen free) animal facilities in The Australian National University or Shandong Analysis and Test Center. C57BL/6 (Jax 000664), CD45.1 (Jax 002014) and *Rag1*^-/-^ (Jax 002216) mice originally from Jackson Laboratory were purchased from Australian Phenomics Facility, Shanghai Model Organisms Center or Vital River Laboratories (VRL). *Il10*^-/-^, *Calca*^-/-^, *Stat1*^-/-^, *Ifnar1*^-/-^ and *Irf1*^-/-^mice were purchased from Shanghai Model Organisms Center. All mice were maintained at a controlled temperature of 23 ± 2 °C, with humidity of 50 ± 10%. All animals had *ad libitum* access to pellet feed and water and were randomly distributed into groups with a maximum of five adult mice per cage. For Ovariectomy (OVX)-induced osteoporosis mouse model, 6-8 weeks old female C57BL/6 wildtype or *Rag1^-/-^* mice were performed ovariectomy (OVX) or sham operation. Mice were subcutaneously injected with 30,000 I.U. recombinant human IL-2 (Beijing SLPharma, Beijing, China) daily 4 weeks after the surgery for another 4 weeks then sacrificed for analysis. Femurs and tibia were collected for analysis. All animal experiment ethics were approved by the designated Animal Ethics Committee of the institutes.

### Microcomputed tomography

To evaluate bone volume and architecture, mice right femurs were fixed in 70% ethanol for μCT imaging. Three-dimensional (3D) scan was applied through the high-resolution μCT scanner (Quantum GX micro-CT Imaging System), and images were acquired at 90 kV voltage and 160 μA tube current, with the integration time of 3 min. A total of 512 binary images were obtained with a wide field of view (FOV), with scanning at 50 mm and a 4.5 μm voxel size resolution. The 3D reconstructions were created by stacking the optimized 2D images from the contoured region. For analysis of trabecular bone, the threshold was set to 75 for analysis of all samples where the acceptable bone architecture could be segregated from the background. The regions of interest of bone were created within the endosteal envelope and were extended 1 mm from the growth plate to proximal metaphysis. The transverse μCT slices were evaluated in a region of interest from 150 slices below the distal growth plate for around 100 consecutive image slices, and the 3D algorithms were used to calculate the relevant parameters, including the bone mineral density (BMD, g/cm3), bone volume fraction (BV/TV, %), trabecular number (Tb.N, 1/mm), trabecular thickness (Tb.Th, mm). All morphometric parameters were analyzed using Quantum GX Analysis Software.

### Bone histology and immunohistochemistry staining

To evaluate the osteoclasts in bone surface, left femurs of mice were collected and fully fixed with 4% paraformaldehyde, paraffin-embedded, cut into 5-μm sections after decalcification. The paraffin embedded sections were stained with H&E or Tartrate-Resistant Acid Phosphatase (TRAP, Servicebio) for the detection of osteoclasts. For immunohistochemistry staining, paraffin embedded sections were blocked with Fc-receptor-blocking antibody (clone 2.4G2, 1:100, BD), followed by staining with TRAP rabbit polyclonal antibody (GB11416, 1:1000, Servicebio). Sections were further washed and stained with secondary antibodies. Slices were permeabilized with xylene and mounted with neutral balsam. All slides were scanned with Pannoramic DESK (3D HISTECH, Hungary). Randomly selected area around proximal metaphysis was analyzed with Pannoramic Viewer (P.V 1.15.3) at the magnification of 10X.

### Flow cytometry

Single cell suspension was washed, counted, and stained with Fc-receptor blocking antibodies (clone 2.4G2, 1:100 dilution, BD) for 10 min on ice to block non-specific staining. For surface staining, cells were washed once with PBS containing 2% heat-inactivated fetal bovine serum (FBS, Gibco) and stained in an appropriately diluted antibody solution for 30 min at 4 °C. 7-amino-actinomycin D (7-AAD) (Biolegend) or Zombie Aqua (Biolegend) were stained to exclude dead cells. For intranuclear staining of transcription factors, cells were washed once after surface staining and fixed using Foxp3/Transcription Factor Staining Buffer Set (eBioscience) for 30 min. The intranuclear specific antibodies were diluted in permeabilization buffer (eBioscience) and incubated for 30 min at room temperature. Data were collected on a BD Aria III or BD LSRFortessa (BD) flow cytometer and analyzed using FlowJo software. Detailed antibody information is in Tables S4.

### Mouse osteoclast differentiation assays

Mouse bone marrow cells were isolated from 6-8 weeks old wild type C57BL/6, *Irf1^-/-^, Stat1^-/-^* and *Ifnar1^-/-^* mice. CD3^-^B220^-^NK1.1^-^CD11b^+^ bone marrow cells were sorted by flow cytometry and differentiated into osteoclasts in the presence of RANKL (50 ng/ml, Biolegend) and M-CSF (20 ng/ml, Peprotech) in 96-well culture plates.

To evaluate CXCR3^+^ T_REG_ cell-mediated regulation of osteoclast formation, bone marrow CD4^+^CD25^+^CXCR3^+^ T_REG_ cells and bone marrow-derived CD3^-^B220^-^NK1.1^-^CD11b^+^ cells were isolated from 6-8-week-old wild type mice. Cells were co-cultured at a ratio of 1:5 (CXCR3^+^ T_REG_/ CD11b^+^) in the presence of of 20 ng/ml M-CSF and 50 ng/ml RANKL for 6 days.

To examine the inhibitory effects by cytokines secreted from ILC2s, flow cytometry sorted ILC2s were ex vivo expanded for 2 days and then rested with or without anti-IL10 or anti-CGRP neutralizing antibodies in α-MEM containing 10% (v/v) FBS for 2 Days. Specifically, ILC2s were sorted as Lineage (CD3, CD5, CD8, CD11b, CD11c, CD19, B220, γδTCR, NK1.1, FcεRIα, TER119, Gr-1)^-^ CD45^+^CD127^+^Sca-1^+^ST2^+^ cells. Culture supernatants were collected and used to treat CD3^-^B220^-^NK1.1^-^CD11b^+^ cells during osteoclastogenesis. To examine ILC2s-mediated regulation of osteoclast formation, flow cytometry sorted CD3^-^B220^-^NK1.1^-^CD11b^+^ cells (6 x 10^4^/well) and freshly cell sorted ILC2s were seed together in different ratios in 96-well plates, in the presence of 20 ng/ml M-CSF and 50 ng/ml RANKL for 5 days.

To examine IL-2-mediated regulation of osteoclast formation, recombinant human IL-2 (Beijing SLPharma, Beijing, China) was added together with M-CSF and RANKL at the indicated concentrations (3000, 1000 or 300 I.U/ml) from day 0.

Osteoclast formation was evaluated by TRAP staining and TRAP^+^ multinucleated cells (nuclei>3) were counted using a brightfield microscope.

### Human peripheral blood mononuclear cells (PBMCs) isolation and osteoclast differentiation

PBMCs were isolated from 20 ml of blood of healthy donors as previously described (*64, 65*). Diluted blood with PBS was then gently loaded to Ficoll-Paque Plus (GE) layer at the ratio of 1:1 followed by density gradient centrifugation (400 g, 20°C, 20 min). Mononuclear cells were collected and processed in PBS for further analysis. For osteoclast formation, human PBMC CD14^+^ cells were purified by FACS sorter and polarized into osteoclastogenesis in the presence of human M-CSF (20 ng/ml) and RANKL (50 ng/ml) in 96 well culture plate for 12 days, with refreshed medium every 2 days. Osteoclast formation was evaluated by TRAP staining and TRAP^+^ multinucleated cells (nuclei > 3) were counted using a brightfield microscope.

### TRAP staining of osteoclasts

Osteoclast formation was examined by staining for TRAP activity. Cells were fixed with 10% formalin for 10 min, permeabilized with ethanol and acetone at a ratio of 1:1 for 1 min at room temperature, and incubated in acetate buffer (pH 5.2) containing naphthol AS-MX phosphate (Sigma) as the substrate and Fast Red Violet LB salt (Sigma) as the dye for the detection of reaction in the presence of 50 mM sodium tartrate. Cells were washed with distilled water and air-dried before analysis using a brightfield microscope.

### Bone resorption assay

Purified mouse bone marrow CD11b^+^ cells were differentiated on dentin slices with 20 ng/ml M-CSF and 50 ng/ml RANKL in the presence or absence of 3000 I.U hIL-2 for 10 days. Cells were removed from the dentin slice by wiping the surface, followed by toluidine blue staining (1g/ml, J.T. Baker, Philipsburg, USA). Images were collected with microscope (NIKON ECLIPSE CI) at the magnification of 40X on randomly selected regions on the slices. The numbers of pit formed by bone resorption on the dentin slices were counted. Image analysis was performed using ImageJ (version 1.54a). Quantification was determined by counting pit area per total area.

### ELISA

The concentrations of IL-10 and CGRP in cultured ILC2 supernatants were measured using mouse IL-10 ELISA Kit (Biolegend) and CGRP ELISA kit (Boster), according to the manufacturer’s instruction. Briefly, assay plates were coated with capture antibodies, respectively, at 4°C overnight. Plates were next washed with PBST (PBS + 0.05% Tween-20) and blocked with bovine serum albumin (BSA) at room temperature for 90 min. Samples from ILC2s culture supernatant were added into the plate with recombinant mouse IL-10 or CGRP used as standards. Next, biotinylated detection antibodies (BioLegend or Boster) were added and incubated at room temperature for 60 min. Avidin-HRP (BioLegend or Boster) conjugates were added and incubated at room temperature for 45 min. Lastly, TMB (Invitrogen) was used for the development and 0.5 M H_2_SO_4_ was used to stop the reaction. Absorbance was detected at the wavelength of 450 nm using a PerkinElmer EnSpire plate reader.

### qPCR

Total RNA was purified from tissue or cells using TRIzol reagent (Invitrogen) or RNeasy Mini kit (Qiagen), and cDNA was synthesized using 5 μg of total RNA with RevertAid first strand cDNA synthesis Kit (Thermo Scientific) then amplified using Fast SYBR^®^ Green Master Mix (Thermo Fisher Scientific) with Applied Biosystem quantitative real-time PCR system. mRNA levels were normalized to β-actin as the endogenous control. The primer sequences used for qRT-PCR were as follows: *Nfatc1*: 5’-CCTCGAACCCTATCGAGTGT-3’ and 5’-CCGATGACTGGGTAGCTG TC-3’; *Acp5*: 5’-GCTGGAAACCATGATCACCT-3’ and 5’-TTGAAGCGCAAACGGTAGTAA -3’; *CathepsinK*: 5’-CTTCCAATACGTGCAGCAGA-3’ and 5’-TCTTCAGGGCTTTCTCGTTC -3’; *Calcr*: 5’-TGCAGACAACTCTTGGTTGG-3’ and 5’-TCGGTTTCTTCTCCTCTGGA-3’; *Dcstamp:* 5’-TTTGCCGCTGTGGACTATCTGC-3’ and 5’-AGACGTGGTTTAGGAATGCAG CTC-3’; *Itgb3*: 5’-CCGGGGGACTTAATGAGACCACTT-3’ and 5’-ACGCCCCAAATCCCACCCATACA-3’; *Il2ra*: 5’-TTAGCATCTGCAAGATGAAG-3’ and 5’-TTCCTCACTAGCCAGA AATC-3’; *Il2rb*: 5’-CTCAAGTGCCACATCCCAGATC-3’ and 5’-AGCACTTCCAGCGGAGA GATCT-3’; *β-actin:* 5’-TGACAGGATGCAGAAGGAGA-3’ and 5’-CGCTCAGGAGGAGCAA TG-3’.

### siRNA

Cultured CD11b^+^ cells were transfected with 40 nM mouse *Il2ra, Il2rb* on-target plus smart pool siRNAs (QIAGEN) using Lipofectamine 2000 (Invitrogen) according to the manufacturer’s instructions. 40 nM negative non-targeting siRNA (QIAGEN) was used as control. 2.5 ml serum-free medium was used for transfection for 24h, and cells were further cultured for 4 days in complete media containing 20 ng/ml M-CSF and 50 ng/ml RANKL for osteoclast formation. After 4-5 days, numbers of osteoclast were examined by TRAP staining.

### Single cell RNA-sequencing

Mouse CD45⁺ bone marrow cells from sham, sham+IL2, OVX, and OVX+IL-2 mice were sorted by flow cytometry for sequencing. 6-8 weeks old female C57BL/6 wildtype mice were used. Cells were loaded onto a Chromium Controller (10x Genomics) for single-cell portioning, followed by library preparation according to the manufacturer’s instructions for the Chromium Next GEM Single Cell 5′ Kit v2. Single-cell libraries and were sequenced on the NovaSeq 6000 (Illumina). All raw data, including read alignment and generation of count matrices, were performed using the Cell Ranger pipeline. Raw base call files generated by sequencing were previously demultiplexed into FASTQ files per sample. The ‘cellranger count’ tool mapped the reads against the murine genome (GRCm38/mm10) and performed unique molecular identifier counting. We utilized a Seurat (v4.1.2) workflow to graphically characterize distinct cell populations and visualize cell clusters. Low-quality cells with <500 detected genes, >6,000 genes, or >5% mitochondrial RNA were removed. The ‘RunHarmony’ function in the Harmony package was employed to remove batch effects in the data set for each sample. Data were normalized and integrated using the ‘SCTransform’ function workflow, followed by PCA for dimensional reduction. For clustering, the ‘FindClusters’ function was used with the resolution set at 0.3 for subsequent analyses. The resultant clusters were visualized using a Uniform Manifold Approximation and Projection (UMAP) dimensionality reduction plot. We employed the ‘FindAllMarkers’ function in Seurat for gene Differential expression analysis, using the Wilcoxon rank sum test with FDR correction, and pathway or module analyses were performed using ‘fgsea’ function and ‘AddModuleScore’ function.

### Bulk RNA-sequencing of CD11b^+^ cells differentiating into osteoclast

Mouse bone marrow CD3^-^B220^-^NK1.1^-^CD11b^+^ cells were sorted by flow cytometry and differentiated into osteoclasts by RANKL (50 ng/ml) and M-CSF (20 ng/ml) with or without IL-2 treatment (3,000 I.U/ml) in 96-well culture plates for 4 days. Total RNA was extracted from the cultured cells using the RNeasy Mini Kit (Qiagen) according to the manufacturer’s instructions. Poly(A)^+^ transcripts were purified, and sequencing libraries with multiplexed barcode adapters were prepared using the NEBNext Ultra II RNA Library Prep Kit for Illumina (NEB) following the manufacturer’s protocol. All samples passed quality control assessment using a Bioanalyzer 2100 (Agilent). High-throughput sequencing (150 bp, paired-end) was performed on an Illumina NovaSeq 6000 platform at Shanghai Biotechnology Corporation, achieving a sequencing depth of 30-50 million reads per sample.

RNA-seq reads were aligned to the murine genome (GRCm38/mm10) using HISAT2 with default parameters. Read counts were obtained using HTSeq-count, and transcript abundances (counts per million, CPM) were calculated with edgeR. Differentially expressed genes (DEGs) were identified using edgeR on three independent biological replicates, with Benjamini-Hochberg false discovery rate (FDR) correction applied for multiple testing. Genes with adjusted p value<0.05 and log fold-change (FC)>0.5 were considered significantly differentially expressed. Pathway enrichment analyses were performed using clusterProfiler and EnrichR. All volcano plots and heat maps were generated using the ggplot2 package in R.

### Bulk RNA-sequencing of ILC2 cells treated with IL-2

RNA-seq data from purified ILC2 treated with IL-2 in Figure 3J were obtained from the Gene Expression Omnibus (GEO; accession number GSE81882) (*26*). FPKM (fragments per kilobase of transcript per million mapped reads) values of the three replicates were defined in the original study. Normalized log2 (FPKM+1) values were used for DEGs identified using limma package together with Benjamini-Hochberg false discovery rate (FDR) correction. Volcano plots were generated using the ggplot2 package in R.

### Data availability

The datasets generated during this study have been deposited into the China National GeneBank Database (CNGBdb) with accession number CNSA: CNP0008767.

### Statistical Analysis

Statistical analyses were performed using GraphPad Prism software (10.0). Comparisons between two groups were assessed using an unpaired, two-tailed Student’s t-test. For comparisons involving more than two groups, a one-way analysis of variance (ANOVA) was performed. Experiments involving two independent variables were analyzed using a two-way ANOVA. A P value < 0.05 was considered statistically significant. Data are presented as mean ± SD or SEM, as indicated in the figure legends.

## Acknowledgments

We thank the patients and healthy individuals for their participation in our study. We thank M. Devoy, H. Vohra, and C. Gillespie of the Flow Cytometry Facility at the John Curtin School of Medical Research for technical support. We further thank the Biomedical Resource Facility at the John Curtin School of Medical Research for assistance with single-cell RNA-sequencing. We thank Shanghai Biotechnology Corporation for bulk-RNA-sequencing.

## Author contributions

P.Z. and T.Z. conceived and oversaw the study. P.Z. and T.Z. designed the experiments. D.Y. participated the initial design of some experiments. P.Z., T.Z., J.F., Y.L. and A.X. performed experiments. Y.Z., D.M. and L.S. collected human data. W.W. provided critical resources. P.Z. and T.Z. analyzed the data and performed the entire bioinformatic analysis. P.Z., T.Z., L.S. and D.Y. supervised the study. P.Z. wrote the manuscript with major help from T.Z. and input from all co-authors.

## Data and materials availability

All data associated with this study are present in the paper or the Supplementary Materials.

