## Supplementary Materials for "Therapeutic interleukin-2 rewires skeletal-immune circuits to reverse postmenopausal bone loss"

**Fig. S1**

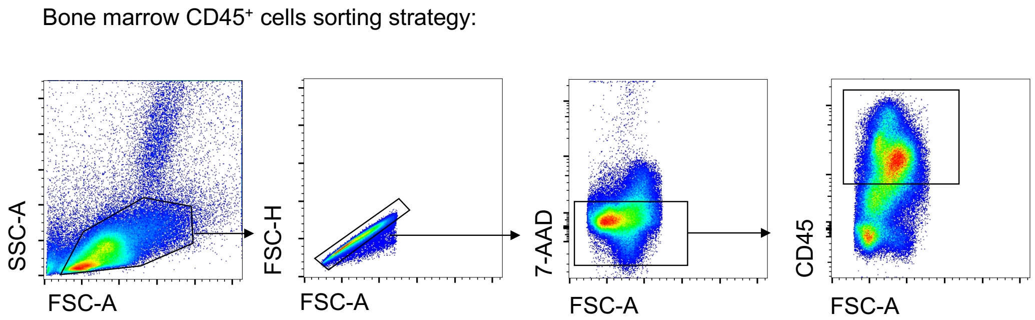


**Fig S1. scRNA-seq analysis of bone marrow CD45⁺ cells.**

Gating strategy for sorting CD45⁺ bone marrow cells used in Fig 1 for single-cell RNA sequencing.

### Fig. S2

**A.**

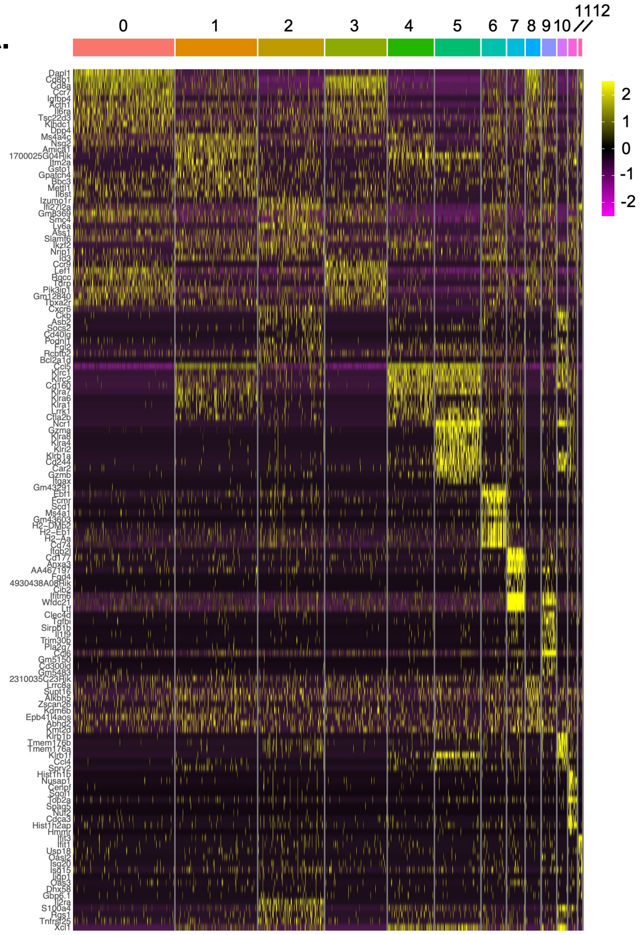


**B.**

Mouse $T_{REG}$ gating strategy

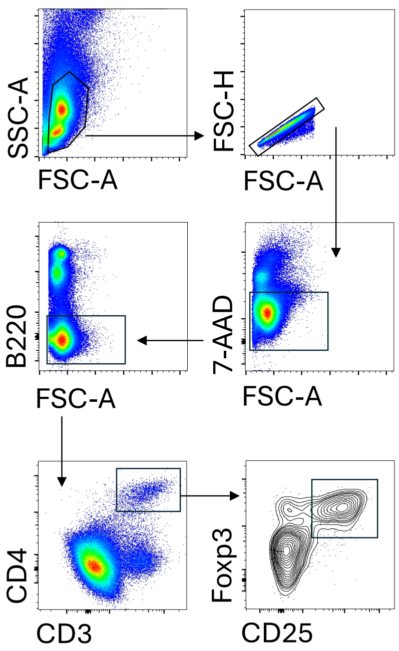


**C.**

Mouse bone marrow CD11b+ gating strategy

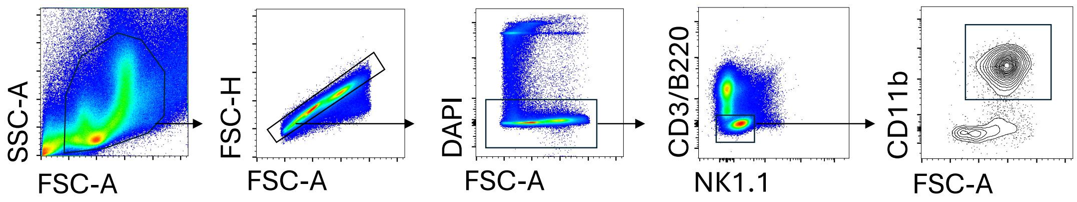


**D.**

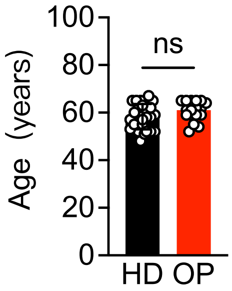

**Fig S2. scRNA-seq analysis of mouse bone marrow $T_{REG}$ cells.**

(**A**) Heatmap showing top 10 genes expression profiles of 13 identified subclusters in bone marrow T cells subsets. (**B**) Gating strategy of mouse bone marrow $T_{REG}$ cells. (**C**) Gating strategy for FACS sorting mouse bone marrow $CD3^-B220^-NK1.1^-CD11b^+$ cells used for *in vitro* osteoclast formation. (**D**) PBMC samples collected from postmenopausal osteoporosis patients (OP) compared to samples from age-matched women healthy donors (HD). Average age of enrolled human participants. T-test, data are means ± SD. ns.: not significant.

**Fig. S3**

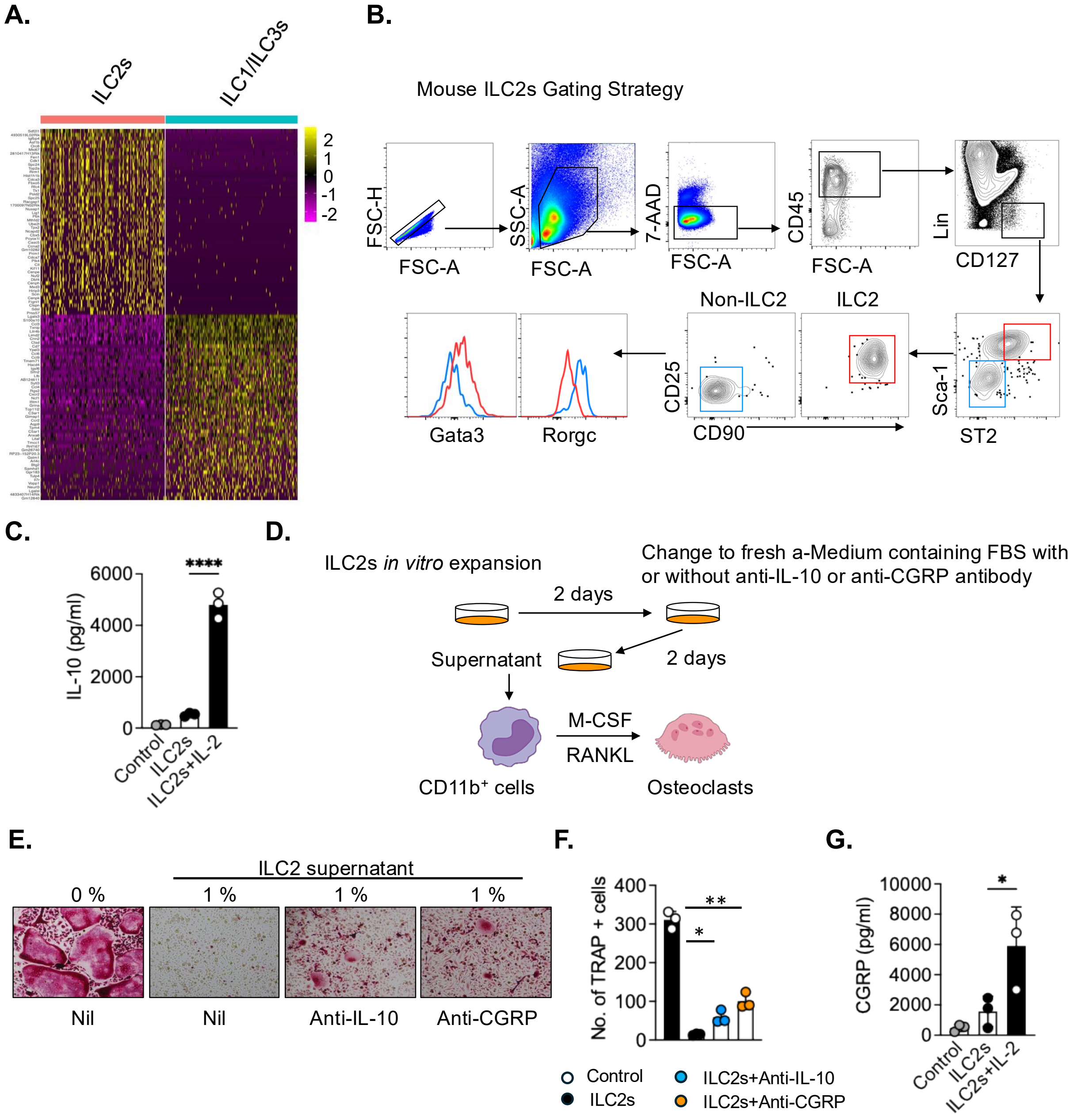


**Fig S3. Characterization of mouse bone marrow ILC2s.**

(**A**) Heatmap showing top 50 genes expression profiles of 2 subclusters in bone marrow innate lymphoid cells (ILCs). (**B**) Gating strategy for the identification of mouse bone marrow ILC2s. (**C**) ELISA of IL-10 production in ILC2 culture supernatants. Representative of at least 2 independent experiments. One-way ANOVA, bar indicates means. ****P<0.0001. (**D**) Schematics of the experiment. The *ex vivo* expanded ILC2s were treated with or without neutralizing antibodies against IL-10 (Anti-IL-10) or CGRP (Anti-CGRP) for 2 days and supernatant were collected and used for treating osteoclastogenesis. $CD3^-B220^-NK1.1^-CD11b^+$ bone marrow cells were differentiated into osteoclasts by 20 ng/ml M-CSF and 50 ng/ml RANKL in the presence or absence of 1% different ILC2-derived culture supernatants for 6 days. Osteoclast formation was evaluated by TRAP staining. (**E**) Representative TRAP staining images. (**F**) Quantification of $TRAP^+$ multinucleated (nuclei > 3) cells. Representative of at least 3 independent experiments. (**G**) ELISA of CGRP production in ILC2s culture supernatants. Representative of at least 2 independent experiments. One-way ANOVA, bar indicates means. $*P<0.05$.

Fig. S4

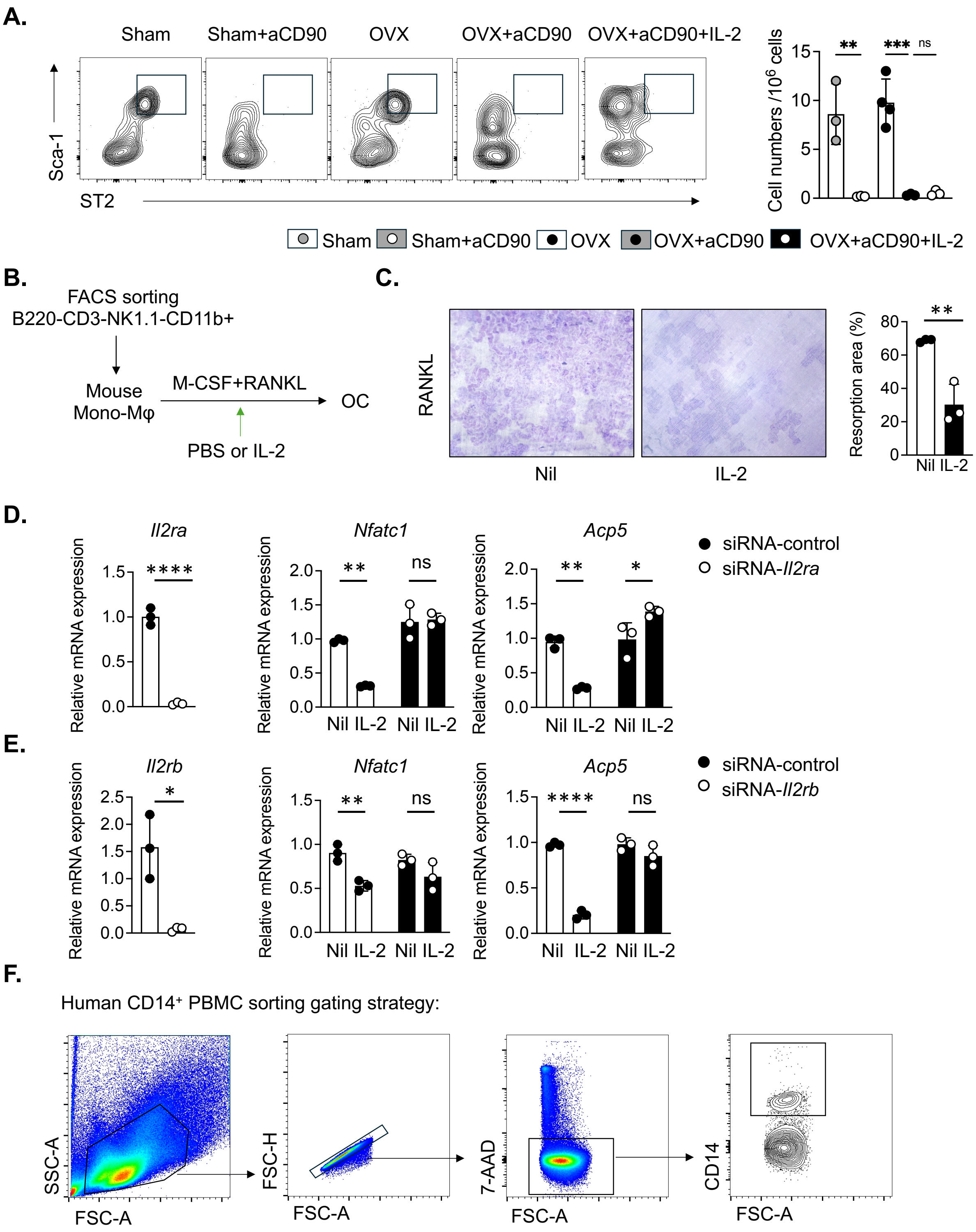
A.
Sham
Sham+aCD90
OVX
OVX+aCD90
OVX+aCD90+IL-2
Sca-1
ST2
Cell numbers /10^6 cells
**
***
ns
Sham
Sham+aCD90
OVX
OVX+aCD90
OVX+aCD90+IL-2
B.
FACS sorting
B220-CD3-NK1.1-CD11b+
Mouse
Mono-Mφ
M-CSF+RANKL
OC
PBS or IL-2
C.
RANKL
Nil
IL-2
Resorption area (%)
**
Nil IL-2
D.
Il2ra
Nfatc1
Acp5
Relative mRNA expression
****
**
ns
*
Nil IL-2 Nil IL-2
siRNA-control
siRNA-Il2ra
E.
Il2rb
Nfatc1
Acp5
Relative mRNA expression
*
**
ns
****
Nil IL-2 Nil IL-2
siRNA-control
siRNA-Il2rb
F.
Human CD14+ PBMC sorting gating strategy:
SSC-A
FSC-A
FSC-H
7-AAD
CD14

**Fig S4. IL-2 directly inhibits osteoclastogenesis.**

(**A**) Anti-CD90 antibody administration in *Rag1*$^{-/-}$ mice to deplete ILCs. Flow cytometry analysis to confirm the efficiency of ILC2 depletion in bone marrow. Representative FACS plots (left) and quantification (right) of BM-ILC2 (Lin$^{-}$CD45$^{+}$CD127$^{+}$ST2$^{+}$Sca-1$^{+}$) in the indicated groups. One-way ANOVA, bar indicates means. ∗∗P<0.01, ∗∗∗P<0.001. (**B**) Schematics of the experiment. Differentiation of FACS sort-purified bone marrow CD3$^{-}$B220$^{-}$NK1.1$^{-}$CD11b$^{+}$ cells into osteoclasts in the presence of RANKL and M-CSF, with 3000 I.U/ml IL-2 added at the start of cell culture. (**C**) CD3$^{-}$B220$^{-}$NK1.1$^{-}$CD11b$^{+}$ cells differentiating into osteoclast on the dentin slices in the presence of with RANKL (50 ng/ml) and M-CSF (20 ng/ml) with or without IL-2 (3,000 I.U/ml) treatment for 10 days. Representative bone resorptive pits staining images (left) and quantification of resorption area (%) by ImageJ. T-test, bar indicates means. ∗∗P<0.01. (**D-E**) Transfection of siRNA targeting *Il2ra* (**D**) or *Il2rb* (**E**) following RANKL-induced osteoclastogenesis of CD3$^{-}$B220$^{-}$NK1.1$^{-}$CD11b$^{+}$ cells with or without IL-2 treatment (3000 I.U/ml). qPCR analysis of relative mRNA expression of the targeted genes (*Il2ra, Il2rb*) or osteoclast marker genes *Nfatc1 and Acp5*. T-test (*Il2a, Il2rb*) or two-way ANOVA (*Nfatc1, Acp5*), bar indicates means. ∗P<0.05, ∗∗P<0.01, ∗∗∗∗P<0.0001. (**A-E**) Representative of at least 2 independent experiments. (**F**) Gating strategy for sorting CD14$^{+}$ human PBMCs used for *in vitro* human osteoclast formation.

**Fig. S5**

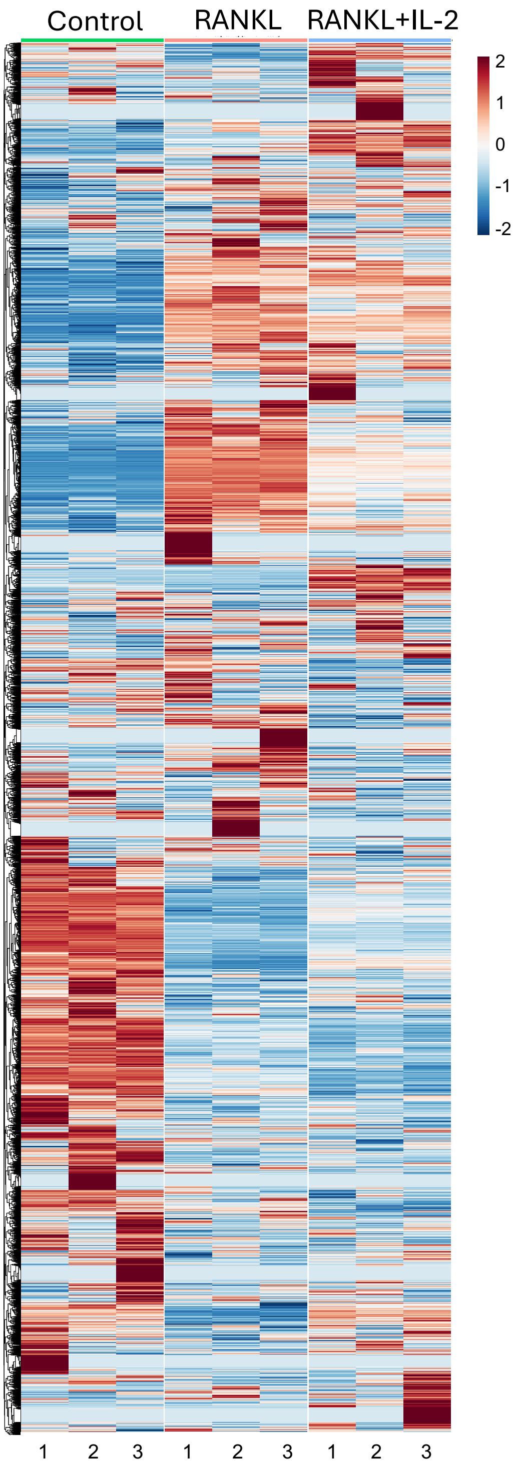


**Fig S5. IL-2 reprograms RANKL-stimulated monocytes.**

Heatmap showing the distinct transcriptional genes profiles across the indicated groups.

**Fig. S6**

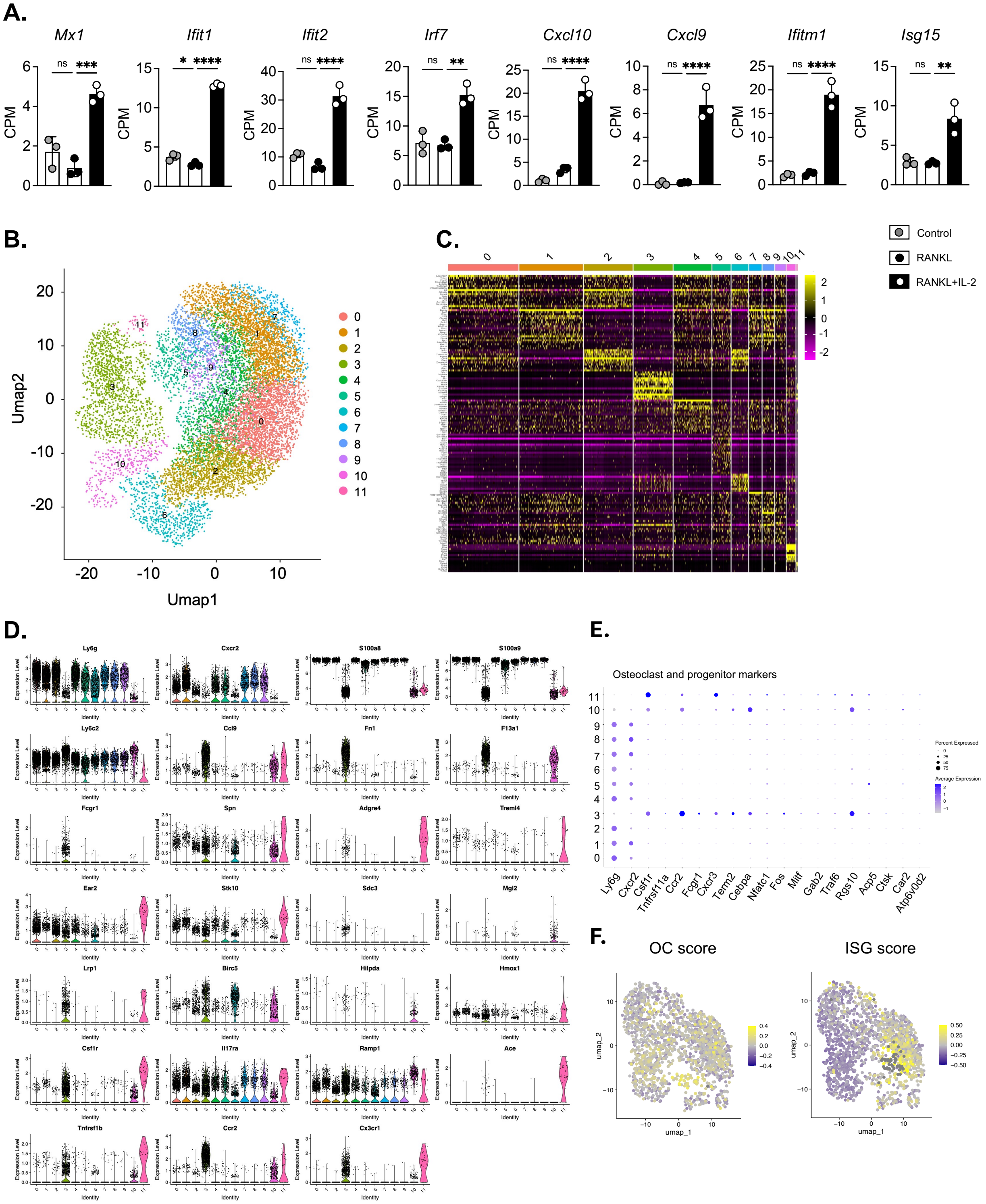
A.
Mx1
Ifit1
Ifit2
Irf7
Cxcl10
Cxcl9
Ifitm1
Isg15
CPM
ns
Control
RANKL
RANKL+IL-2
B.
Umap2
Umap1
C.
D.
Ly6g
Cxcr2
S100a8
S100a9
Ly6c2
Ccl9
Fn1
F13a1
Fcgr1
Spn
Adgre4
Treml4
Ear2
Stk10
Sdc3
Mgl2
Lrp1
Birc5
Hilpda
Hmox1
Csf1r
Il17ra
Ramp1
Ace
Tnfrsf1b
Ccr2
Cx3cr1
Expression Level
Identity
E.
Osteoclast and progenitor markers
Ly6g
Cxcr2
Csf1r
Tnfrsf11a
Ccr2
Fcgr1
Cxcr3
Trem2
Cebpa
Nfatc1
Fos
Mitf
Gab2
Traf6
Rgs10
Acp5
Ctsk
Car2
Atp6v0d2
Percent Expressed
Average Expression
F.
OC score
ISG score
umap_2
umap_1

**Fig S6. Activation of IFN-program is essential for the IL-2-mediated intrinsic suppression of osteoclastogenesis.**

(**A**) Normalized expression (CPM) of representative interferon-stimulated genes (ISGs) in the indicated groups. One-way ANOVA, bar indicates means. ∗∗P<0.01, ∗∗∗P<0.001, ∗∗∗∗P<0.0001. (**B**) UMAP visualization of bone marrow myeloid cells subsets (cluster 1, 2, 6 from **Fig.1F**). (**C**) Heatmap of the top 20 differentially expressed genes (DEGs) defining each of the 12 subclusters. (**D**) Violin plots showing genes associated with osteoclast or progenitors across the 12 subclusters. (**E**) Dot plot of classical marker genes of osteoclast or progenitors (*Tnfrsf11a, Ctsk, Acp5, Nfatc1*, etc.) in 12 identified clusters. (**F**) Feature plots showing the osteoclast (OC) and interferon stimulating genes (ISG) module scores within OC progenitor-enriched clusters. Osteoclast related gene list used for OC score: *Ca2, Calcr, Dcstamp, Ocstamp, Acp5, Mmp9, Jdp2, Car2, Fos, Dnmt3a, Acot1, Cpt2, Mfn1, Prkar2a, Serpinf1*. Interferon stimulating gene list used for ISG score: *Mx1, Ifit1, Cxcl10, Isg15, Oas1a, Ifit3, Ifit2, Irf7, Oasl2, Samd9l, Parp9, Herc6, Igtp, Ifi47, Ddx58.*

**Table S1. Clinical characteristics of healthy donor and osteoporosis patients.**

| | Healthy Donor (n=27) | Osteoporosis (n=17) | p-value |
|---|---|---|---|
| **Gender** | Female | Female | / |
| **Age (yr)** | 58.3 (48-67) | 60.7 (52-65) | 0.13 |
| **BMI (kg/m$^2$)** | 24.4 (21.2-31.25) | 24.8 (22.6-28.12) | 0.4867 |
| **Cholesterol (mmol/L)** | 5.134 (4.05-7.31) | 5.246 (3.6-6.87) | 0.7344 |
| **Triglyceride (mmol)** | 1.725 (0.74-3.45) | 1.587 (1.09-2.83) | 0.6301 |
| **Low density lipoprotein (LDL)** | 2.797 (1.77-4.65) | 3.28 (2.24-4.65) | 0.0768 |
| **High density lipoprotein (HDL)** | 1.516 (0.9-3.13) | 1.271 (0.85-1.59) | 0.1005 |
| **Urea nitrogen (mmol/L)** | 5.032 (2.7-7.67) | 4.684 (3.32-5.2) | 0.4848 |
| **T-score** | -0.4182 (-2.4-2.9) | -3.508 (-4.0--3.0) | <0.0001 |
| **Z-score** | 0.4727 (-1.8-3.5) | -1.915 (-2.8--1.3) | <0.0001 |

**Table S2. Gene list used for enrichment analysis of CXCR3$^+$ Treg signature.**

| **Gene list used for enrichment analysis of CXCR3$^+$ Treg signature** |
|---|
| Ccl5, Cxcr3, Nkg7, Ifng, Tbx21, AW112010, Igtp, Stat1, Il2rb, Epsti1, Dusp2, Gimap4, Izumo1r, Rac2, Nr4a2, Irf1, Gimap3, Ifi47, Aplp2, AU020206, Gimap7, Cbx4, Gzmb, Bhlhe40, Gbp7, Wipf1, Agfg1, Gimap6, Pglyrp1, Ccr5, Icos, Gnas, Samhd1, Maf, Nfatc1, Itgb2, Malat1, Tap1, Nedd9, Ptpn22, Pla2g16, Psme1, Cebpb, Vps37b, Ctla4, Sorl1, Mbnl1, Cd3e, Rabgap1l, Ypel3, Lbh, Psmb8, Lck, Dusp5, Irf2bpl, Itgal, Smc4, Ikzf2, Cdkn1b, Trac, Cbl, Clk1, Itpkb, Trbc2, Cd2, Snhg5, Ifi203, Tln1, Gm8995, Rsrp1, Ly6a |

**Table S3. Gene list used for enrichment analysis of *Stat5b*-CA signature.**

| **Gene list used for enrichment analysis of *Stat5b*-CA signature** |
|---|
| Cish, Socs2, Klrg1, Il1rl1, Il9r, Tnfsf8, F2r, Ptafr, Itgb3, Cd47, Selplg, Itgb7, Lama5, Lgals1, Drc1, Myo6, Frmd5, Actr3b, Vim, Rhoc, Il1r2, Tox, Cxcr6, Cd200, Cxcl10, Irf8, Stat5b |

**Table S4. Antibody list.**

| Mouse Antibodies | | | |
|---|---|---|---|
| **Antigen** | **Conjugates** | **Clone** | **Vendor** |
| **CD11b** | FITC | M1/70 | BD |
| **CD90.2** | FITC | 30-H12 | BD |
| **7AAD** | / | / | Thermo Fisher |
| **CD25** | PE | PC61 | BD |
| **GATA3** | PE-eFlour 610 | TWAJ | eBioscience |
| **PE-CF594 streptavidin** | PE-CF594 | / | BD |
| **Sca-1** | PE-cy7 | D7 | Biolegend |
| **APC streptavidin** | APC | / | BD |
| **Foxp3** | AF647 | 150D | Biolegend |
| **CD3** | Alexa Fluor 700 | 17A2 | Biolegend |
| **CD45** | APC-cy7 | 30-F11 | BD |
| **B220** | APC-cy7 | RA3-6B2 | BD |
| **CD122** | Brilliant Violet 421 | TM-β1 | BD |
| **ST2** | Brilliant Violet 421 | DIH9 | Biolegend |
| **CD11b** | Brilliant Violet 510 | M1/70 | BD |
| **Zombie Aqua** | Brilliant Violet 510 | / | Thermo Fisher |
| **CD4** | Brilliant Violet 605 | RM4-5 | BD |
| **CD127** | Brilliant Violet 605 | A7R34 | Biolegend |
| **CD3** | Brilliant Violet 650 | 17A2 | Biolegend |
| **CD25** | Brilliant Violet 650 | PC61 | Biolegend |
| **Sca-1** | Brilliant Violet 786 | D7 | Biolegend |
| **CD3** | Biotin | 17A2 | Biolegend |
| **CD4** | Biotin | GK1.5 | Biolegend |
| **CD19** | Biotin | 6D5 | Biolegend |
| **B220** | Biotin | RA3-6B2 | Biolegend |
| **NK1.1** | Biotin | PK136 | Biolegend |
| **CD11c** | Biotin | N418 | Biolegend |
| **CD11b** | Biotin | M1/70 | Biolegend |
| **Ly6G/Ly6C（Gr-1)** | Biotin | RB6-8C5 | Biolegend |
| **gdTCR** | Biotin | GL3 | Biolegend |
| **TER119** | Biotin | TER-119 | Biolegend |
| **CD5** | Biotin | 53-7.3 | BD |
| **CD8a** | Biotin | 53-6.7 | BD |
| **FceRIα** | Biotin | MAR-1 | Biolegend |
| **Human Antibodies** | | | |
| **CD45** | FITC | HI30 | Biolegend |

| | | | |
|---|---|---|---|
| **CRTH2** | FITC | BM16 | Biolegend |
| **TCRab** | FITC | IP26 | Biolegend |
| **CCR6** | PE | 29-2L17 | Biolegend |
| **7AAD** | / | / | Thermo Fisher |
| **CD161** | PE | HP-3G10 | Biolegend |
| **CD25** | PE-CF594 | M-A251 | BD |
| **PE-CF594 streptavidin** | PE-CF594 | / | BD |
| **CD127** | BV421 | A019D5 | Biolegend |
| **APC streptavidin** | APC | / | BD |
| **CD4** | AF700 | RPA-T4 | BD |
| **CD45** | AF700 | HI30 | Biolegend |
| **CD45RA** | APC-Cy7 | HI100 | BD |
| **CXCR3** | BV421 | G025H7 | Biolegend |
| **CCR6** | BV510 | G034E3 | Biolegend |
| **CD127** | BV605 | A019D5 | Biolegend |
| **CD3** | BV650 | SK7 | BD |
| **CD25** | BV650 | BC96 | Biolegend |
| **CD14** | BV786 | M5E2 | Biolegend |
| **CD19** | BV786 | SJ25C1 | BD |
| **CD56** | APC-Cy7 | HCD56 | Biolegend |
| **CD3** | Biotin | UCHT1 | Biolegend |
| **CD4** | Biotin | RPA-T4 | Biolegend |
| **CD19** | Biotin | HIB19 | Biolegend |
| **CD16** | Biotin | 3G8 | Biolegend |
| **CD14** | Biotin | 63D3 | Biolegend |
| **CD19** | BV786 | SJ25C1 | BD |
| **CD3** | Biotin | UCHT1 | Biolegend |
| **CD4** | Biotin | RPA-T4 | Biolegend |
| **CD19** | Biotin | HIB19 | Biolegend |
| **CD16** | Biotin | 3G8 | Biolegend |
| **CD14** | Biotin | 63D3 | Biolegend |
| **CD20** | Biotin | 2H7 | Biolegend |
| **CD34** | Biotin | 581 | Biolegend |
| **CD123** | Biotin | 6H6 | Biolegend |
| **TCRalpha/beta** | Biotin | IP26 | Biolegend |
| **HLA-DR** | Biotin | L243 | Biolegend |
| **FcεRIα** | Biotin | AER-37 (CRA-1) | Biolegend |
| **CD56** | Biotin | HCD56 | Biolegend |
